# IPSC-based modeling of resiliency in centenarians reveals longevity-specific signatures

**DOI:** 10.1101/2025.10.10.681757

**Authors:** Todd W. Dowrey, Samuel F. Cranston, Nicholas Skvir, Yvonne Lok, Payton Bock, Elizabeth K. Kharitonova, Elise MacDonald, Ella Zeldich, Christopher Gabel, Alexander Tyshkovskiy, Stefano Monti, Vadim N. Gladyshev, Paola Sebastiani, Thomas T. Perls, Stacy L. Andersen, George J. Murphy

## Abstract

Centenarians represent a human model of resilience to age-related decline, yet resiliency mechanisms remain elusive. Here, we establish an induced pluripotent stem cell (iPSC)-based platform to interrogate resilience signatures in centenarians. IPSC-derived neurons from centenarians exhibit transcriptional programs promoting synaptic integrity, calcium homeostasis, and cholesterol biosynthesis, while suppressing proteostatic stress pathways. Functionally, these neurons maintain stable calcium dynamics, reduced baseline mitochondrial activity, and energy-efficient homeostasis. Upon challenge, centenarian-derived neurons mount a robust stress response, in contrast to attenuated responses in non-centenarian controls. This resilience signature parallels adaptations in long-lived mammals and aligns with healthy brain aging, while showing erosion in Alzheimer’s disease and cancer. Our platform provides a scalable human model for dissecting resilience biology offering a framework to extend healthspan and mitigate age-related decline.

## INTRODUCTION

The human population has been steadily growing, reaching an estimated 8.2 billion people as of 2024(*1*). In the United States, the increase in individuals over 70 years of age has outpaced that of those under 70 since 1950(*2*). More simply put, the human population is growing disproportionately older. This trend has put a significant financial and logistic burden on the healthcare system to treat an aging population, as age is the main risk factor for many chronic diseases like Alzheimer’s disease (AD), cerebrovascular and cardiovascular disease(*3*, *4*). Understanding the factors and mechanisms that promote healthy aging is critical if we are to develop therapies and strategies that effectively compress morbidity and disability towards the very end of longer and healthier lifespans.

Throughout life, humans are constantly exposed to varying forms of insult, such as infection or stress. “Resilience” is our ability to respond to these insults, mount an effective cellular response, and restore normal physiological function(*5*, *6*). However, resiliency declines with age, hindering the ability to restore healthy function after insult. Along these lines, molecular entropy increases with age and is implicated in the onset of many hallmarks of aging and related disease(*7*, *8*). A potential underlying cause of aging-associated decline in resilience is entropy(*9*). For example, increased entropy at the level of DNA methylation predicts mortality in BXD mice(*10*). This often coincides with an increase in frailty, or loss of functional capacity, and onset of disability and chronic disease. Frailty can be identified by functional deficits such as muscle loss or fatigue(*11*), and chronic diseases are diagnosable with known symptomology and clinical biomarkers(*12–14*). Just as molecular mechanisms may drive disease and disarray, we hypothesize that understanding and identifying the underpinnings of resilience against or resistance to decline may allow for the delay or prevention of frailty and chronic disease. Moreover, these resiliency mechanisms would provide promising targets for therapeutics that may slow or reverse aging.

Centenarians, those who live over 100 years, are natural human models of extended healthspan, or the years of life lived without disability, disease, or cognitive impairment(*15*). Centenarians display resilience or resistance to heart disease, cancer, Alzheimer’s disease and related disorders (ADRD)(*16–20*) and the ability to compress morbidity(*21*) much later into life than the typical ager (*18–20*, *22–24*). Interestingly, many centenarians and their offspring display significantly lower biological age compared to their chronological age using molecular aging clocks thus suggesting the existence of molecular drivers of healthful aging and longevity(*25*, *26*). Previous work by our group and others has characterized the centenarian immune system and shown a unique maintenance of immune functionality and enrichment for longevity-associated factors even after a history of exposure to insult(*27*, *28*). Given this, we believe that EL subjects provide the blueprint for understanding how to improve functionality and cellular performance, ultimately delaying or eliminating aging-related functional decline and disease.

Induced pluripotent stem cells (iPSCs) have emerged as a powerful model system that act as a renewable source of biomaterial for the modeling of complex diseases and the preclinical screening of therapeutics(*29*, *30*). We have generated a first-of-its-kind bank of iPSCs from EL subjects to fuel the modeling of resiliency (Dowrey et al. 2024). These cell lines capture the genetics of EL subjects which allows for the investigation of the documented heritability of EL(*31–33*) as well as specific longevity-associated gene variants and genetic signatures (*34–39*). Importantly, iPSC-based models have been applied to study diseases of aging which take decades to manifest, such as neurodegenerative disease(*40–45*) and amyloid disease(*46–48*). Here, we present a novel iPSC-based model of resiliency in which cell types of aging-related interest, such as forebrain neurons, are generated from EL and non-EL subjects to identify the molecular and functional signatures of exceptional brain aging and perhaps more generally, exceptional longevity. We discovered a molecular and functional resiliency signature consisting of genes involved in neuronal development and stress response as well as functional enhancement of both energetics and neuronal signaling regulation. Produced cells were then exposed to cellular stress in an effort to understand how stress response signatures differ between EL and non-EL subjects after insult. We found that EL subjects who demonstrated lower levels of entropy prior to insult, showed an enhanced molecular response to stress. Together, these results highlight a distinct molecular and functional signature in EL subjects that improves cellular regulation and performance. Our results also provide insight into how these signatures may be used to understand aging-related disease trajectories as well as develop novel therapeutics that may slow or reverse neurological aging.

## RESULTS

### Application of diverse biological aging clock models allows for selection of EL subjects at the extremes of resiliency

To fuel the establishment of an iPSC-based model of resiliency, centenarian subjects identified to be at the extremes of resiliency at the time of iPSC blood draw were selected as detailed in Dowrey et al(*49*). Briefly, subjects were stratified through clinical history, functional profiling (Barthel Activity of Daily Living (ADL) Index(*50*)), and cognitive assessment data. In combination with these assessments and as a molecular register of relative health for the subjects that would be carried forward for stem cell-based modeling in this study, we performed DNA methylation profiling on the source peripheral blood mononuclear cells (PBMCs) and employed an array of biological aging clocks. These clocks included the GrimAge clock(*51*), PC-based updates to the Horvath, PhenoAge, and Hannum DNA methylation-based clocks(*52*), as well as the recently developed intrinsic capacity (ICAge)(*53*) and AdaptAge clocks(*54*). The models include principal component-based versions of four classic and well-established models (which reduce technical variance relative to the original versions(*52*)), as well as two recently published modern models (IC Age and AdaptAge).

To briefly summarize these models, each is based on average methylation states at a number of specific sites across the genome, derived from training predictive models using chronological age, as well as various clinical and biological information depending on the model. The Horvath clock, trained solely on chronological age and outputting predicted age in years, was one of the original pan-tissue age predictors that established epigenetic clocks as a viable metric by which to assess biological age, and is often used as a benchmark readout(*55*). The Hannum clock is trained in the same way but specifically with blood as opposed to multiple tissues(*7*). PhenoAge was trained on chronological age, as well as multiple clinical mortality risk markers, serving as a better predictor of mortality risk and healthspan(*56*). GrimAge was developed by training on long-term survival data to most effectively estimate direct mortality risk(*51*). The Intrinsic Capacity (IC) clock was trained on a composite score reflecting functional reserve and resilience (locomotion, cognition, vitality, sensory, and psychological well-being) and reflects changes in overall functional capacity with age(*53*). Lastly, AdaptAge was one of a series of models trained using CpGs showing causal links to aging traits – the CpGs from AdaptAge are correlate specifically to ‘adaptive’ or more protective health outcomes with age(*54*). Thus, its age-acceleration (relative to chronological age) is interpreted to represent the degree of beneficial/adaptive response.

Across the clocks which estimate biological age (PC_Horvath, PC_Hannum, and GrimAge), EL subjects displayed reduced biological age compared to their chronological age (**Fig. 1A-C**). Similarly, EL subjects showed a spectrum of health and phenotype predictions in the PC_PhenoAge, ICAge, and AdaptAge clocks (**Fig. 1D-F**) which may allow for further subdivision of groups in future studies to investigate the correlation between phenotypic readouts and disease risk. Overall, we employed a combinatorial strategy of assessing clinical, functional, and molecular readouts when choosing exceptional longevity (EL) subjects to inform a novel iPSC-based model of human resiliency.

**Fig. 1.**
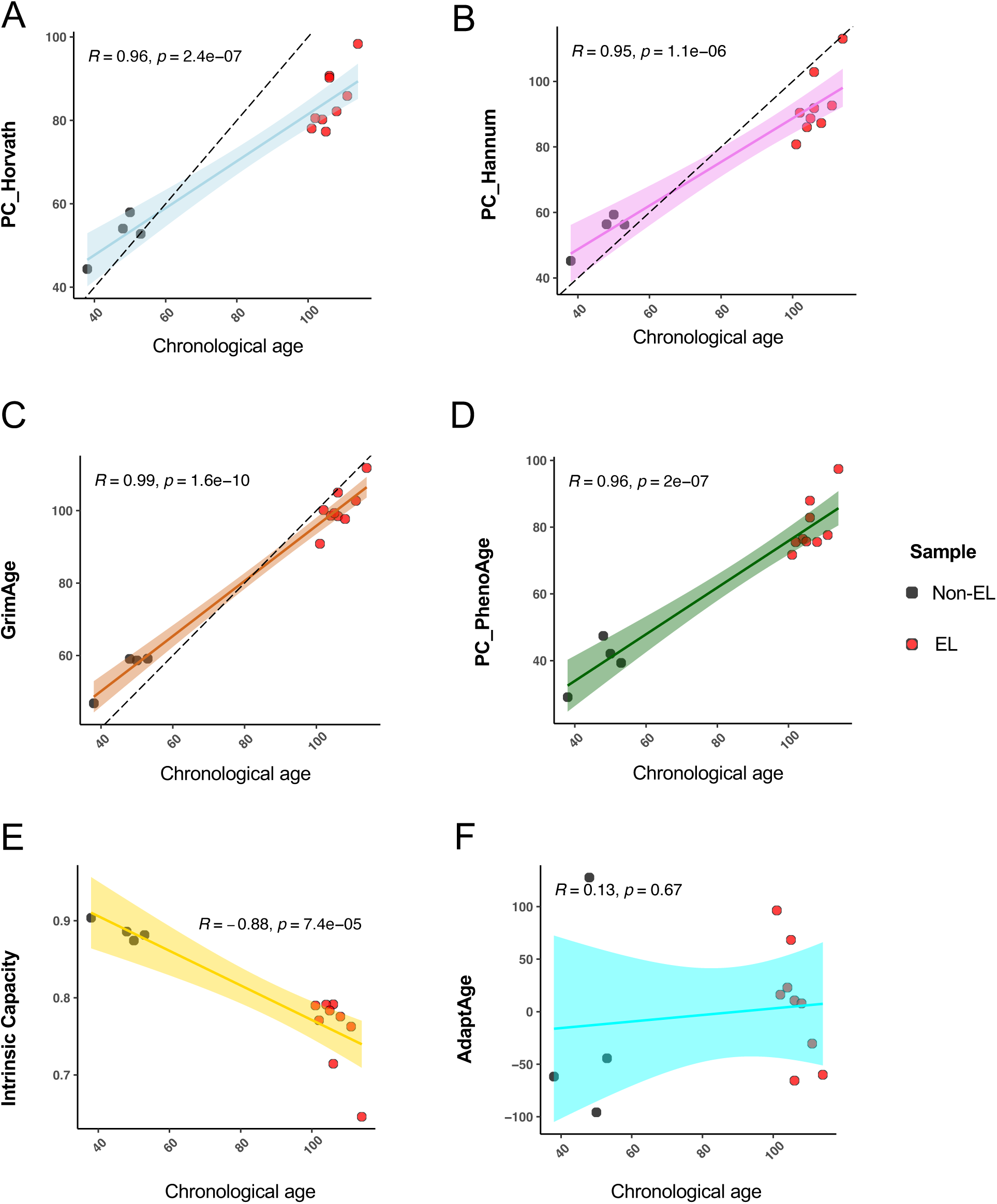
Centenarians display reduced biological age based on a multitude of aging clocks. Biological age and related estimates for EL (red points) and non-EL (green points) peripheral blood mononuclear cells analyzed using the PC_Horvath (**A**), PC_Hannum (**B**), GrimAge (**C**), PC_PhenoAge (**D**), Intrinsic Capacity (**E**), and AdaptAge (**F**) biological clocks. The colored lines represent the best linear fit from each model, while the dotted black lines show a theoretical linear correlation between chronological and biological age, where applicable.

### Centenarian iPSC-derived neurons display molecular signatures of resilience

To investigate exceptional longevity in the context of neuronal resilience, we generated iPSC-derived cortical forebrain neurons from centenarian (EL) subjects as well as non-EL controls without family history of longevity. Neuronal differentiation was achieved using a NGN2-mediated forward programming strategy(*49*) (**Fig. 2A, Fig. S1**). Under baseline, unperturbed conditions, bulk RNA sequencing of produced cells revealed a distinct transcriptional signature in EL-derived neurons (**Fig. 2**). Notably, EL neurons showed significant upregulation of genes implicated in calcium signaling, synaptic integrity, and neuronal maturation, including *Neurocalcin delta* (NCALD) and *MVK*, which encodes for Mevalonate kinase(*57–59*). NCALD is a neuronal calcium-binding protein that buffers intracellular calcium levels(*60*, *61*). Mevalonate kinase is involved in cholesterol biosynthesis(*62*), a critical determinant of membrane structure, synaptic function, and neuronal signaling as well as mitochondrial dynamics (**Fig. 2B**)(*63–66*). The enrichment of these pathways suggests that EL neurons may sustain enhanced calcium trafficking, membrane properties, and synaptic integrity.

**Fig. 2.**
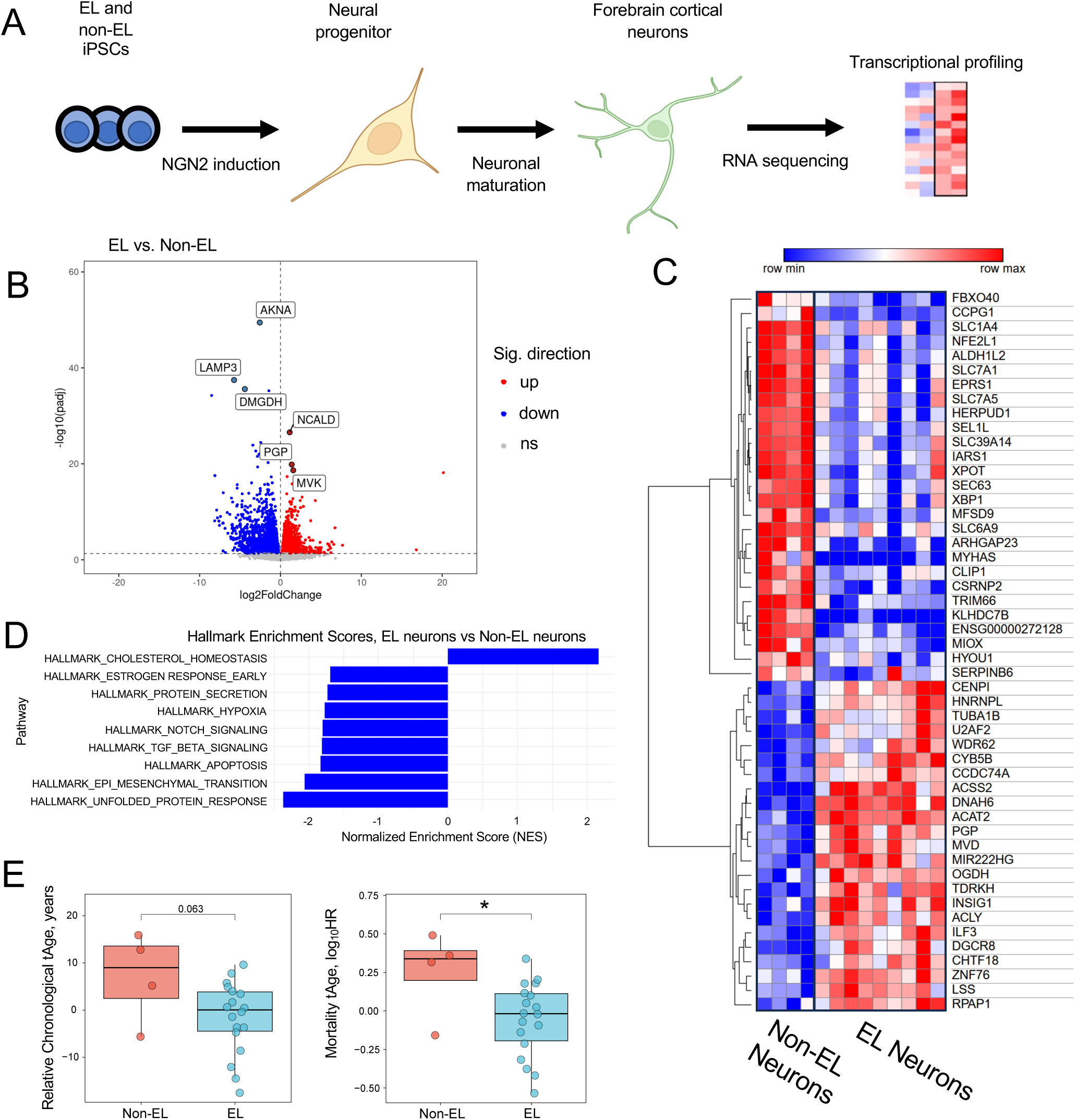
Centenarian iPSC-derived neurons display unique, longevity-associated molecular signatures. **A**) Experimental outline detailing the production of forebrain cortical neurons from EL and non-EL iPSC lines for transcriptional profiling. **B**) Volcano plot comparing gene expression between EL and Non-EL neurons (Red = significantly upregulated in EL, blue = significantly downregulated, gray = not significant). **C**) Heatmap displaying top 50 differentially expressed genes (DEGs) between EL and non-EL neurons (Red = upregulated, Blue = downregulated). **D**) Hallmark pathway enrichment based on DEGs from EL neuron vs non-EL neuron comparison (Blue = significantly enriched pathway (p>0.05)). **E**) Multispecies conventional Chronological tAge (left) and mortality tAge (right) predictions of EL and Non-EL neurons (paired statistical test, *=p<0.05).

In contrast, non-EL neurons displayed upregulation of stress response genes responding to proteostasis disruption and activation of the unfolded protein response (UPR), including HERPUD1(*67*), HYOU1(*68*), and XBP1(*69*, *70*) (**Fig. 2C–D**). This pattern indicates that, even at baseline conditions, non-EL neurons experience greater proteotoxic stress and a heightened demand for cellular stress-response mechanisms.

To further evaluate differences in biological age between the groups, we applied transcription-based, multi-tissue aging clocks of chronological age and expected mortality(*71*) to iPSC-derived neurons. EL-derived neurons exhibited a significantly reduced mortality transcriptional age (tAge) compared to non-EL neurons according to the mortality clock and a consistent trend toward reduced chronological tAge (p=0.063) according to the chronological clock (**Fig. 2E, Fig. S2**).

Altogether, these findings indicate that neurons derived from centenarian iPSCs are characterized by enhanced expression of genes that promote synaptic health and neuronal resilience, whereas non-EL neurons exhibit signatures of proteostasis strain. These results suggest that EL neurons have improved regulation of cellular functions and evade molecular hallmarks of aging, such as loss of proteostasis(*72*).

### Live calcium imaging reveals improved regulation of neuronal activity in centenarian neurons

To assess whether transcriptional resilience signatures observed in EL-derived neurons translate into distinct functional dynamics, we performed live-cell calcium imaging. iPSC-derived cortical neurons from EL and non-EL backgrounds were transduced with the fluorescent calcium reporter GCaMP8s (**See methods**), and somatic calcium events were quantified (**Fig. 3A-C**). Aligning with transcriptional differences, EL neurons exhibited a marked decrease in the percentage of active neurons compared to non-EL neurons. Additionally, EL neurons displayed significantly shorter individual calcium events than non-EL neurons (**Fig. 3D**). Notably, the event widths of EL-derived neurons are more consistent (coefficient of variance 0.469) compared to non-EL (coefficient of variance 0.744). However, the interpeak interval (IPI) does not differ within active neurons in the two data sets (**Fig. 3F**). This combination of molecular (**Fig. 2**) and functional data supports the hypothesis that EL-derived neurons intrinsically maintain more balanced neuronal dynamics and system-wide activity compared to non-EL neurons.

**Fig. 3.**
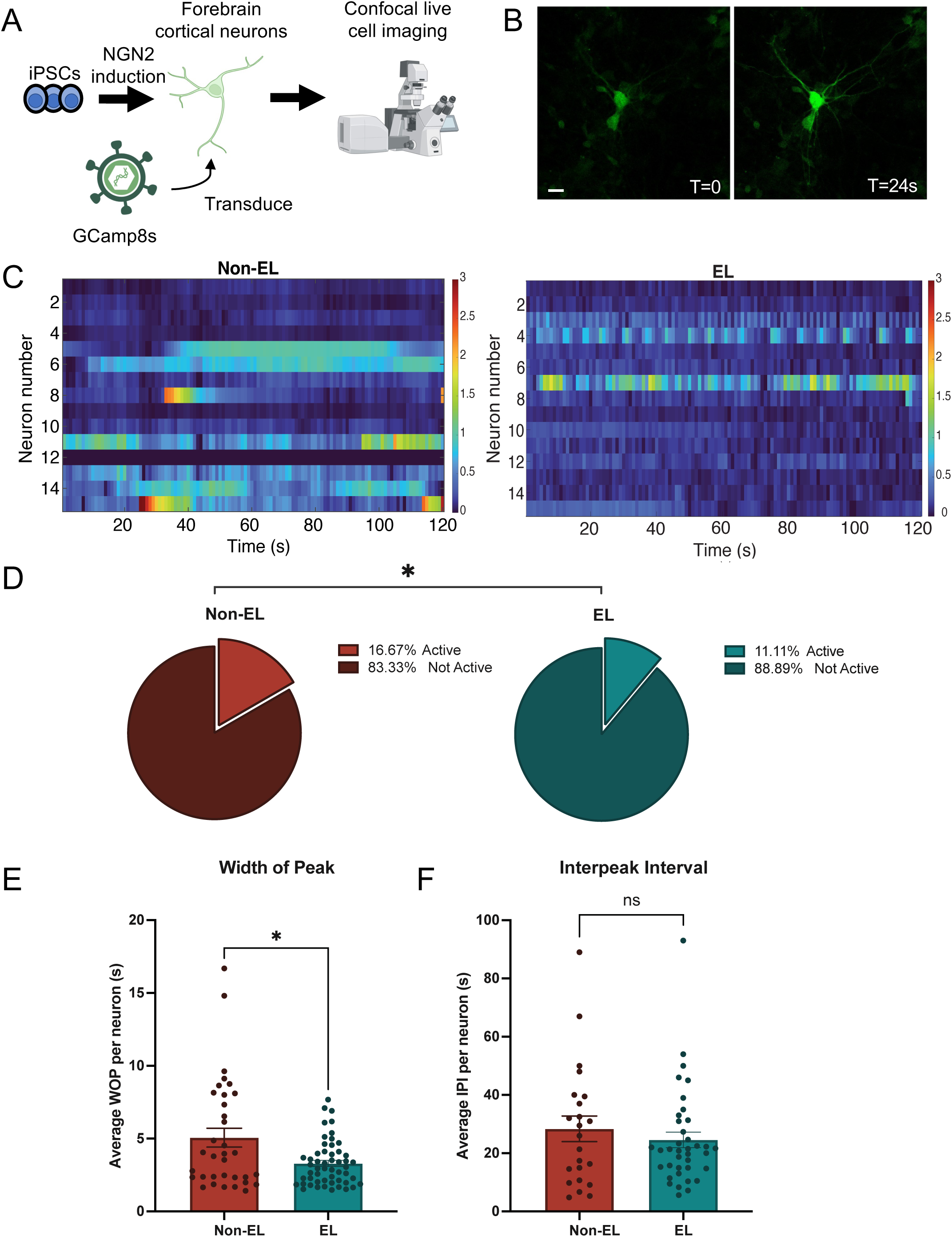
Live calcium imaging reveals improved regulation of neuronal calcium dynamics and synaptic transmission in EL neurons. **A**) Experimental outline detailing the production of forebrain cortical neurons from EL and non-EL iPSC lines, transduction with fluorescent calcium reporter, and live cell confocal imaging. **B**) Confocal images of calcium spike (Scale bar=20µm, movie in supplement). **C**) Heatmap representation of neuronal calcium activity where each row is one ROI (soma) imaged across 120 seconds (x axis) (Red represents higher intensity, blue represents baseline). **D**) Percent activity of total neurons imaged across 4 non-EL iPSC lines (red) and 9 EL iPSC lines (blue)(*=p<0.05). **E**) Bar plot of average width of calcium signal peak where each dot represents a neuron across non-EL (Red) and EL (Blue) groups. **F**) Bar plot of average interpeak interval (time between peaks) where each dot represents a neuron across non-EL (Red) and EL (Blue) groups.

### Centenarian-derived neurons display reduced mitochondrial membrane potential and activity

To explore whether resilience in EL-derived neurons extends to mitochondrial regulation and cellular energetics, we assessed mitochondrial dynamics using MitoTracker™ dyes. Total mitochondrial content was quantified with MitoGreen (MTG), while polarized mitochondria were measured with MitoRed (MTR). The ratio of MTR/MTG, reflecting mitochondrial activity and membrane potential, was visualized using confocal microscopy and quantified via flow cytometry (**Fig. 4A–D**). Under baseline, unperturbed conditions, EL-derived neurons exhibited a lower mitochondrial membrane potential and reduced activity compared to non-EL controls (**Fig. 4C-D**). This suggests that EL neurons operate in a more energy-efficient state, avoiding mitochondrial hyperactivity at rest.

**Fig. 4.**
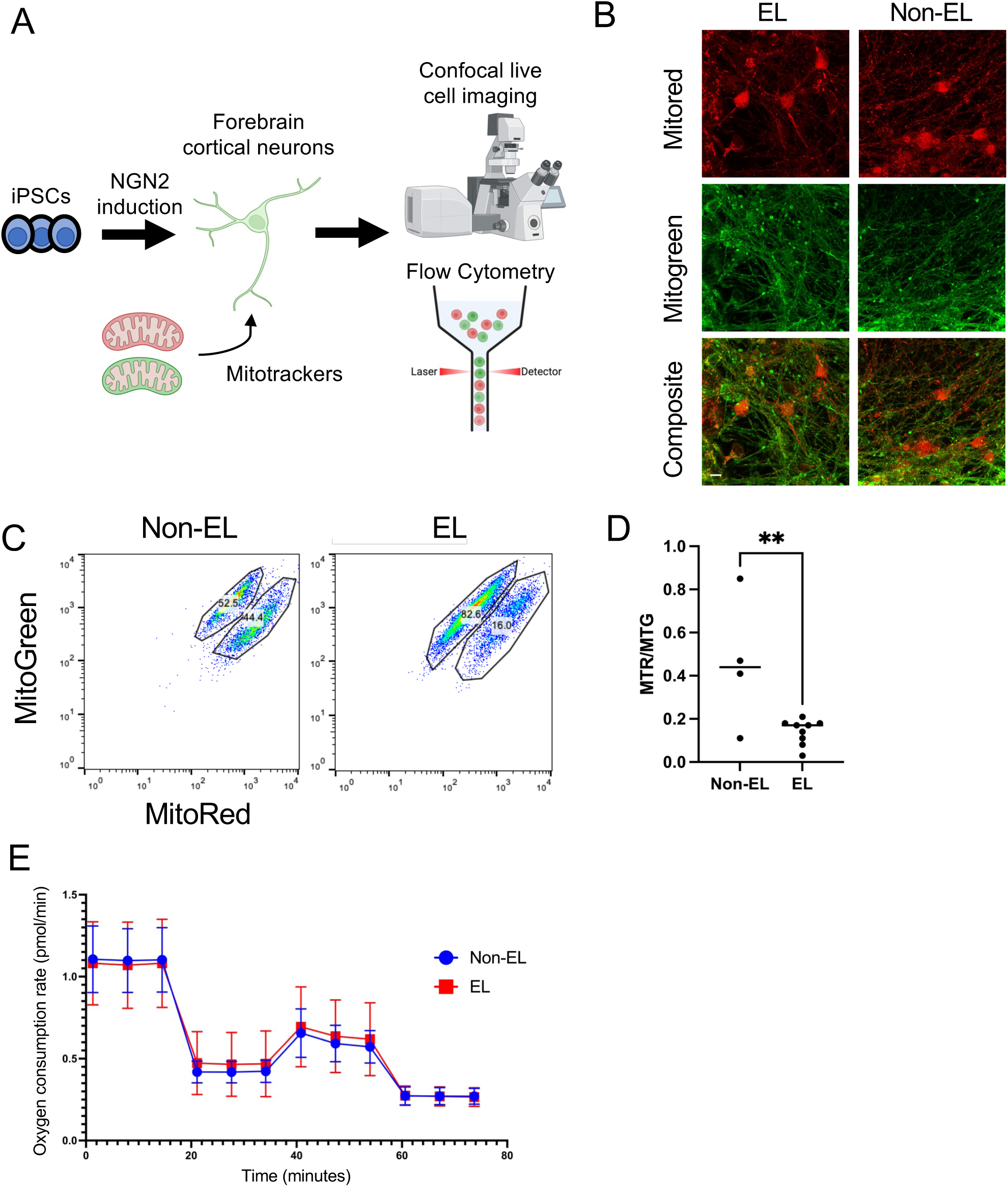
EL neurons exhibit reduced mitochondrial activity and membrane potential. **A**) Experimental outline detailing the production of forebrain cortical neurons from EL and non-EL iPSC lines, transduction with fluorescent calcium reporter, and live cell confocal imaging or flow cytometry. **B**) Confocal imaging of neurons stained with MitoTrackers red and green (scale bar=10µm). **C**) Flow cytometry of cell populations following MitoTracker staining with upper population positive for MitoGreen and lower population positive for both MitoGreen and MitoRed. **D**) Quantification of 4 non-EL and 9 EL cell lines in profiled in flow cytometry MitoTracker experiments (p<0.01). **E**) Seahorse Mito Stress Test displaying oxygen consumption rate across EL and non-EL groups.

To test whether these differences altered responses to metabolic stress, we performed a Seahorse mitochondrial stress assay. Surprisingly, no significant differences were observed between EL and non-EL neurons in stress-induced mitochondrial function (**Fig. 4E**). These findings indicate that, despite having a reduced basal mitochondrial activity, EL neurons retain normal adaptive responses to metabolic stress. Overall, these results suggest that centenarian-derived neurons maintain lower mitochondrial activity at baseline, potentially decreasing oxidative damage(*73*, *74*), while preserving the capacity to respond effectively under stress.

### Centenarian-derived neurons exhibit dynamic resilience to proteostatic stress

Given the baseline enhancements of EL neurons in maintaining proteostasis (**Fig. 2**), we next examined their ability to respond to acute proteostatic disruption. IPSC-derived neurons from EL and non-EL donors were exposed to thapsigargin(*47*, *75*, *76*) a potent ER stressor that disrupts protein folding and secretion, and resultant transcriptional responses were profiled by RNA sequencing (**Fig. 5, Fig. S3**). Interestingly, genes associated with the unfolded protein response (UPR) and stress response genes that respond to proteostasis disruption—many of which were enriched in non-EL neurons at baseline—were significantly upregulated in EL neurons compared to non-EL neurons following stress. For instance, AKNA and LAMP3, genes involved in neurogenesis, neuroinflammation, and UPR regulation(*77–79*), were significantly upregulated in EL neurons post-thapsigargin exposure (**Fig. 5B–C**) compared to non-EL neurons.

**Fig. 5.**
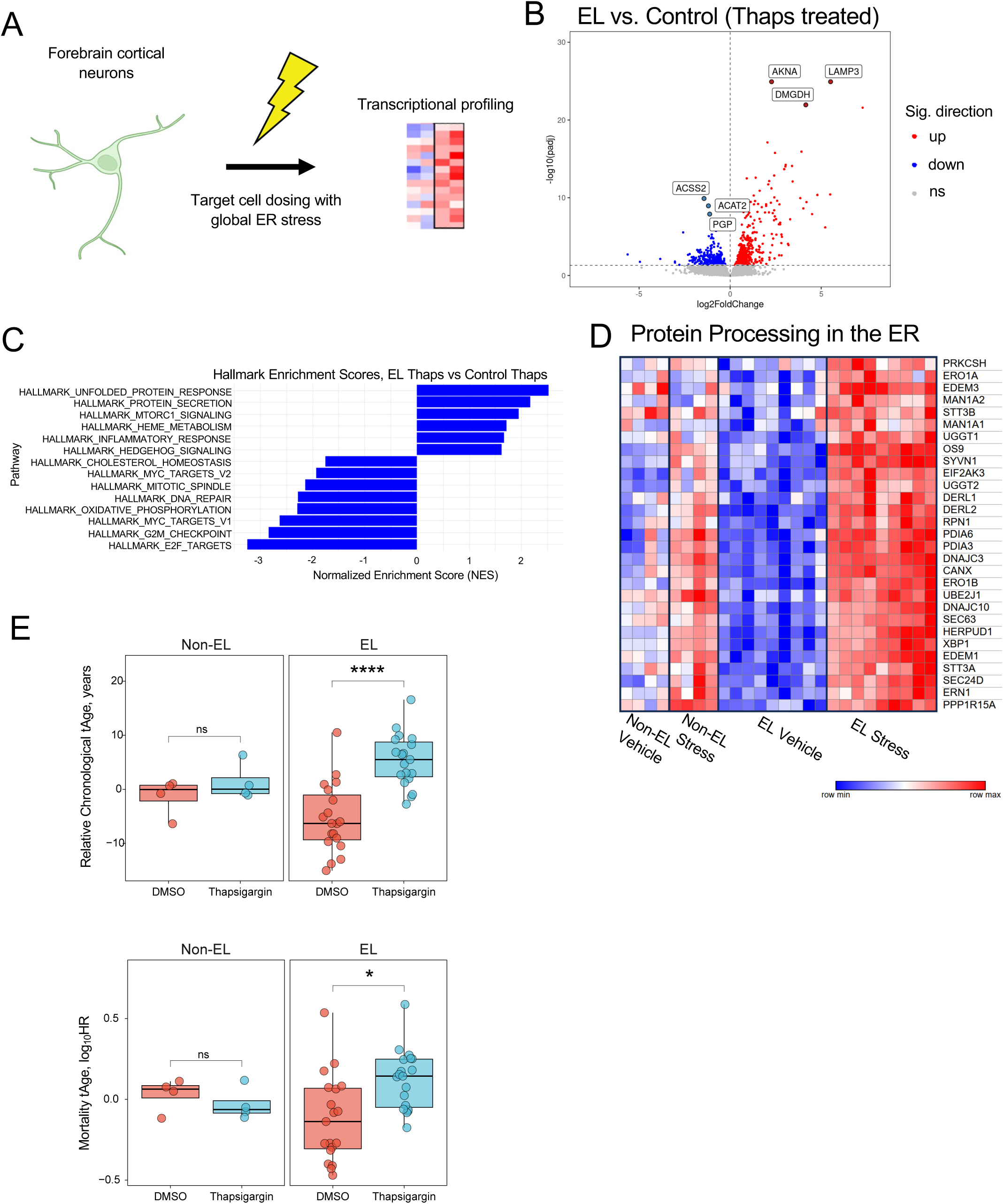
A longevity-specific stress response signature in EL iPSC-derived neurons reveals enrichment for resiliency factors and stress response pathways. **A**) Experimental outline detailing the production of forebrain cortical neurons from EL and non-EL iPSC lines and subsequent application of cellular stress for transcriptional and functional profiling. **B**) Volcano plot comparing gene expression between EL and control neurons following stress (Red = significantly upregulated, blue – significantly downregulated, gray (not significant). **C**) Normalized pathway enrichment score based on overall gene expression differences between EL and non-EL neurons following thapsigargin exposure. **D**) Protein processing in the ER (KEGG) gene set showing expression values across each subject with and without stress. **E**) Multispecies conventional Chronological tAge (top) and mortality tAge (bottom) predictions of EL and Non-EL neurons treated with DMSO (Red) or Thapsigargin (Blue). (paired statistical test adjusting for patient ID, *=p<0.05, ****=p<0.0001).

Pathway analysis further revealed that the top EL-enriched KEGG pathway, Protein Processing in the ER, showed strikingly different dynamics between the EL and non-EL groups. Prior to stress, these genes displayed chronic low-level (“smoldering”) expression in non-EL neurons, suggestive of ongoing proteostasis disruption. In contrast, EL neurons maintained little to no expression at baseline but mounted a robust and coordinated upregulation of this pathway only after thapsigargin challenge (**Fig. 5D**). Lastly, we applied transcriptional aging clocks to assess how stress impacted cellular biological age. Non-EL neurons showed little to no change in chronological or mortality tAge following stress exposure, consistent with their already elevated baseline stress signature. By contrast, EL neurons displayed a significant increase according to both clocks (**Fig. 5E**), reflecting their dynamic activation of stress response programs rather than chronic activation at rest.

Collectively, these findings illustrate a critical distinction: non-EL neurons exist in a state of chronic loss of proteostasis, blunting their capacity to respond dynamically to extrinsic proteostatic stress, whereas EL neurons maintain a more stable baseline and deploy robust, coordinated responses only when challenged. This stress-sensitivity defines a key axis of resilience in centenarian-derived neurons.

### Longevity-specific stress response signatures as a predictive model of resilience to age-related disease

To determine whether the resilience signature we identified in centenarian-derived neurons is unique to our model or has broader applicability, we compared our results against transcriptional datasets from age-related diseases. First, we tested the EL neuronal stress-response signature against post-mortem brain tissue from late-onset Alzheimer’s disease (LOAD) and age-matched healthy subjects(*80*) (**Fig. 5**). The EL resilience signature was significantly more similar to healthy aging brains than to LOAD brains, with the strongest overlap in the dorsolateral prefrontal cortex and visual cortex (**Fig. 6A–C**). This suggests that loss of EL-like resilience pathways may be a feature of Alzheimer’s disease, particularly in cortical regions.

**Fig. 6.**
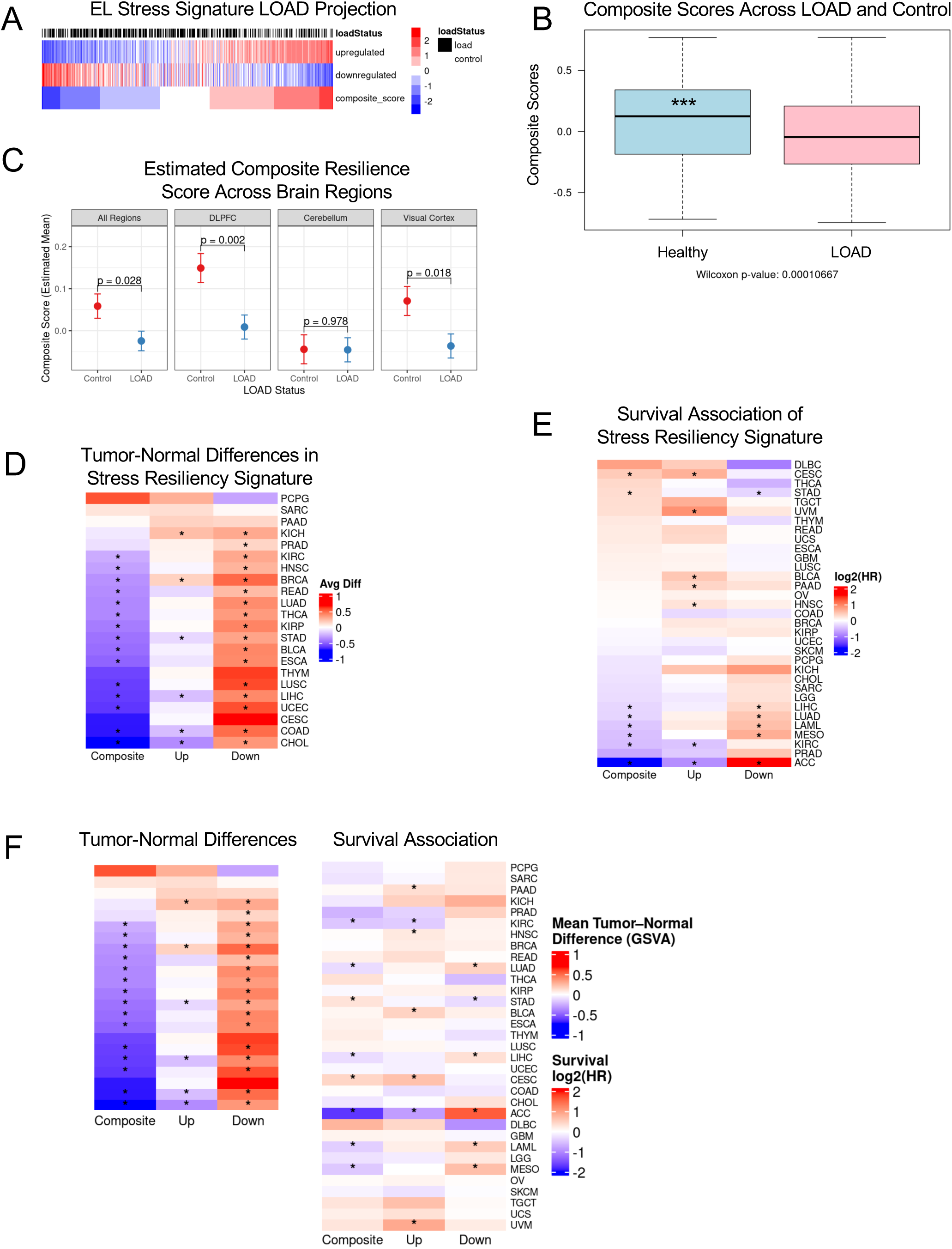
EL stress resilience signature as a predictive model of age-related disease risk and cancer outcomes **A)** Projection of the EL-stress signature onto the LOAD cohort (GSVA heatmap). Rows show GSVA scores for the EL-stress associated upregulated, downregulated, and composite (Up–Down) gene sets; columns are individual LOAD samples, ordered by the composite score (low → high). **B)** Estimated marginal mean GSVA composite scores (Up – Down) from a linear mixed-effect model in control versus LOAD samples (top). Composite resilience score of EL resiliency signature to healthy (Blue) or LOAD (Red) subjects (bottom). **C**) Composite resilience score across brain regions in healthy (Red) or LOAD (Blue) subjects. **D**) Heatmap of average GSVA score differences (Tumor − Normal) for the EL-stress resiliency signature across cancer types, shown separately for the composite score, upregulated and downregulated gene sets. Rows represent cancer types (ordered by composite difference), and columns represent signature components. Colors indicate the direction and magnitude of tumor–normal differences (blue = lower in tumor, red = higher in tumor). Asterisks mark cancer–signature combinations with nominal *p* < 0.05 (Wilcoxon signed-rank test). **E**) Heatmap of log₂ hazard ratios (HRs) from Cox proportional hazards models relating the EL-stress resiliency signature to overall survival, shown for the composite score, upregulated and downregulated gene sets. Rows represent cancer types (ordered by the composite HR), and columns represent signature components. Colors indicate the direction and magnitude of survival associations (blue = lower risk, red = higher risk). Asterisks mark cancer–signature combinations with nominal *p* < 0.05. **F**) Paired heatmaps display tumor– normal differences (left) and survival associations (right) for the EL-stress resiliency signature. Rows represent cancer types (ordered by the mean composite tumor–normal difference), and columns represent the composite, upregulated, and downregulated gene set scores. In the tumor– normal panel, colors indicate mean GSVA score differences between tumors and matched normal tissues (blue = lower in tumors, red = higher in tumors), with asterisks marking cancer–signature combinations significant at nominal *p* < 0.05 (Wilcoxon signed-rank test). In the survival panel, colors indicate log₂ hazard ratios (HRs) from Cox proportional hazards models relating each signature score to overall survival (blue = protective, red = higher risk), with asterisks marking associations with nominal *p* < 0.05. Grey indicates cancers with unavailable normal data.

We next asked whether the EL resilience signature could also provide insights into cancer biology and associated risk. When compared to 33 TCGA (The Cancer Genome Atlas) cohorts(*81*), the EL signature showed significant associations with overall survival in 13 cancer types. Interestingly, the direction of association varied. In some cancers, higher EL signature activity correlated with better survival, while in others it correlated with poorer outcomes (**Fig. 6D-E**). These divergent patterns suggest that resilience pathways may interact with tumor biology in context-dependent ways, shaping disease aggressiveness and prognosis. Finally, we examined how EL-associated features change during carcinogenesis by comparing matched tumor and adjacent normal tissues across 22 cancer types. In 15 of these cancers, tumors showed a significant reduction in the EL resilience score, driven largely by loss of genes that were downregulated in the EL stress response (**Fig. 6F**). Overall, tumors exhibited systematically lower EL scores than expected by chance, indicating that cancer progression may be broadly associated with erosion of resilience pathways. These analyses demonstrate that the transcriptional resilience signature observed in EL neurons may extend beyond our *in vitro* model. Its presence in healthy aging brain tissue, loss in Alzheimer’s disease, and variable influence across cancers all highlight the clinical relevance of resilience pathways as potential predictors of disease susceptibility and outcomes.

## Discussion

### A Multilayered Signature of Neuronal Resilience

Our analyses reveal a consistent theme of resilience in neurons derived from centenarian iPSCs. At the molecular level, EL neurons preferentially upregulate genes that support synaptic integrity, calcium signaling, and cholesterol biosynthesis, while avoiding heightened activation of proteostatic stress response pathways observed in non-EL controls (**Fig. 2**). At the functional level, this transcriptional signature translates into improved regulation of neuronal performance. Live-cell calcium imaging demonstrated that EL neurons exhibit more regular calcium dynamics with shorter peak widths and longer inter-peak intervals, consistent with a quieter, more balanced functional state. In contrast and relative to EL neurons, non-EL neurons showed persistent baseline dysregulation more common with disease states such as epilepsy(*82*), paralleling their stress-associated gene expression profiles (**Fig. 2-3**). At the metabolic level, EL neurons displayed reduced mitochondrial membrane potential and basal activity (**Fig. 4**). Notably, this finding was independent of cellular ER stress (**Fig S4**). This energy-efficient state likely limits oxidative stress(*83*, *84*) while maintaining the ability to mount normal adaptive responses under metabolic challenge, as evidenced by equivalent performance to non-EL cells in Seahorse stress assays.

Once this system was disrupted with extrinsic insult, here achieved through thapsigargin-induced ER stress, EL neurons displayed dynamic resilience demonstrated by upregulation of genes such as AKNA and LAMP3 and pathway enrichment including genes associated with protein processing in the ER. Non-EL neurons, however, showed chronic, low-level activation of proteostasis stress response genes prior to stress, and failed to mount as effective of a response as EL neurons following insult (**Fig. 5**). Further, transcriptional aging clocks reflected these differences in adaptability. EL neurons showed significant increases in tAge following stress, while non-EL neurons exhibited little change, consistent with muted responsiveness (**Fig. 5E**).

These patterns converge on a unified model of human resiliency. In our assays, centenarian-derived neurons are uniquely molecularly programmed, functionally tuned, metabolically efficient, and dynamically adaptable. They are less burdened by chronic proteostasis stress, maintain quieter and more controlled neuronal signaling, conserve energy through reduced mitochondrial activity, and retain the capacity to mount strong, coordinated responses under acute challenge. In contrast, non-EL neurons exhibit early hallmarks of vulnerability, including chronic stress activation, functional dysregulation, and diminished adaptive flexibility.

### Conserved resilience strategies in long-lived mammals: parallels with the bowhead whale and naked mole rat

Aspects of the multilayered resilience signature we observe in centenarian-derived neurons, including reduced baseline mitochondrial activity, lower proteostatic stress, and robust, compensatory stress responses, closely parallels adaptations described in other exceptionally long-lived mammals. The bowhead whale (*Balaena mysticetus*), which can live over 200 years, is a long-lived mammal whose molecular profile aligns well with that of centenarian-derived neurons. Transcriptomic analyses of the bowhead whale reveal alterations in energy regulation, including reduced activity of IGF/insulin pathway modulators such as Grb14, consistent with lower basal metabolic drive and enhanced energy efficiency(*85*). These changes mirror our observation that EL neurons maintain reduced mitochondrial membrane potential at baseline, a state that likely limits oxidative damage while preserving stress responsiveness. In addition, bowhead whale cells demonstrate enhanced DNA repair capacity and attenuated chronic stress signaling(*85*), echoing the way EL neurons avoid baseline proteostatic activation yet mount a coordinated unfolded protein response (UPR) when challenged. Together, these features highlight a shared strategy of conserving energy and minimizing damage during steady state, while preserving robust adaptive capacity when stress arises.

Additionally, long-lived rodent outliers such as the naked mole rat (NMR)(*86–88*) demonstrate a similar longevity strategy seen in centenarians. The NMR maintains exceptional proteostasis through multiple mechanisms. For example, ultra-high-molecular-mass hyaluronan, a contact inhibition polysaccharide speculated to have evolved higher concentrations in the NMR to deal with tunnel dwelling, contributes to their exceptional cancer resistance(*89*). Moreover, the NMR displays increased translational fidelity(*90*) that limits misfolded proteins and elevated proteasome activity(*91*, *92*), which inherently lower basal damage while preserving the ability to clear insults efficiently. These mechanisms resonate with the lower mitochondrial baseline activity and reduced proteostasis burden, coupled with rapid transcriptional activation under stress that is specifically noted in EL neurons.

### Human in vitro models to discover and validate geroprotectors

The underlying mechanisms that drive resiliency or, conversely, rapid decline, remain unclear. Moreover, models of *human* aging, longevity, and resilience to aging-associated disease that allow for the functional testing of potential interventions are virtually non-existent(*93–95*). To directly address these gaps in understanding, our human, iPSC-based model of resiliency provides a system in which to identify and validate biomarkers of aging and perform preclinical screening of novel therapeutics aimed at improving resiliency and extending healthspan. Importantly, this model is malleable and can be easily adapted and scaled to study other biological questions of aging and resiliency by changing the cell or tissue type as well as the form of insult or stress introduced to the system. Further, this model allows for a variety of additional control cohorts to be included, such as those from accelerated aging disorders(*96–99*), to investigate molecular and functional differences across aging rates.

### Limitations of the study

Here, we propose a new model of human resiliency by employing iPSC-based models to study centenarian exceptional longevity. As a byproduct of the reprogramming process, there is a resetting of the epigenetic landscape of the parent cell. In the context of aging-related models, this presents a limitation in investigating the epigenetic changes that are accrued throughout life. However, we believe this platform presents a valuable opportunity to study how and what epigenetic changes result from life processes such as development and exposure to insult in a systematic process. Moreover, iPSCs allow for the study of genetic background which is implicated as a driver of epigenetic changes IPSC-based models provide a powerful human system in which to study disease processes and development(*100*). A main advantage of these models is the flexibility and scalability to produce any cell type of the body to fuel these studies. In this publication, we provide a proof-of-concept centered in the brain – a body system of high interest in the context of aging and neurodegeneration – through the patterning of iPSCs to cortical forebrain neurons. Even so, we have performed epigenetic profiling of the source peripheral blood cells from the centenarians in our bank published previously(*49*) and expanded upon in this work to gain insights into the epigenetic landscape of these subjects at the time of collection. Further, a limitation is that we focus on studying resilience only in the context of cortical forebrain neurons, a cell type strongly impacted by aging. However, given the inherent flexibility of iPSC-based systems, this model is easily adaptable through the tailoring of cell type and perturbation used to new research questions.

## Acknowledgments

We acknowledge all study participants for making this work possible. ChatGPT was used as an editing tool for clarity.

## Funding

This work was supported by

NIH/NIA AG064704

NIH/NIA AG023122

NIH/NIA AG073172

McKnight Brain Research Foundation

## Author contributions

Conceptualization: TWD, TTP, SLA GJM,

Methodology: TWD, SFC, NS, YL, PB, EKK, EM, EZ, CG, AT, SM, VNG, GJM

Investigation: TWD, SFC, NS, YL, PB, EKK, EM, EZ, CG, AT, SM, VNG, GJM

Visualization: TWD, NS, PB, EKK, EZ, CG, AT, SM Funding acquisition: PS, TTP, SLA, GJM

Writing – original draft: TWD, NS, PB, EKK, AT, GJM

Writing – review & editing: TWD, NS, PB, EKK, EZ, CG, AT, PS, TTP, GJM;

## Data and materials availability

Data and materials are available through the ELITE portal (https://elite portal.synapse.org/).

## Supplementary Materials

### Materials and Methods

#### PBMC characterization using biological aging clocks

DNA was extracted from isolated PBMCs using the Qiagen DNeasy Blood and Tissue kit (Qiagen, 69506) according to manufacturer protocols. DNA methylation profiling was performed using the Illumina HumanMethylationEPIC v2 BeadChip array. Initial quality control was performed using the Minfi package (v1.54.1)(*101*) to compute mean detection p-values for each CpG site and sample, filtering those with mean detection values >0.05 from further analysis. Normalization of signal intensities was performed using single-sample Noob normalization (ssNoob) via the preprocessNoob function in Minfi, using the “single” dye bias correction method. Beta values were further collapsed to probe family identifiers using the betasCollapseToPfx function from the Sesame R package (v1.26.0)(*102*). Missing CpGs were imputed using reference data from the sesameData package. Following quality control and normalization, the resulting beta values were used as input for the estimation of epigenetic age across a panel established methylation clocks. Final predictions were returned as either estimated biological ages in years or unitless age-associated scores, depending on the design of each model.

#### IPSC creation and utilization

IPSC lines used in this study were generated as described in our previous publication, Dowrey et al. 2024. Briefly, collected and isolated PBMCs were reprogrammed using a sendai virus-mediated delivery of the OSKM factors per manufacturers protocol (Fisher Scientific, A16517). Resultant clones were expanded, validated, and expanded for study.

#### Generation of iPSC-derived forebrain cortical neurons

Cortical forebrain neurons were generated using a forward programming methodology as described in Dowrey et al. 2024. Briefly, cells were engineered to have a doxycycline-inducible neurogenin 2 (NGN2) cassette allowing for homogenous generation of neural progenitor cells (NPCs) in 3 days of induction. Following induction, NPCs were replated onto a poly-d-lysine, poly-L-ornithine, Laminin coated matrix for further maturation with growth factors for 14 days. At day 17, cells were used in the assays described.

#### Bulk RNA sequencing

RNA was isolated from cells using the Qiagen Allprep DNA/RNA isolation kit (Qiagen, 80284). Bulk RNA sequencing was performed by Novogene Co. (Sacramento, CA) using the Illumina Novaseq6000 S4 Flowcell paired-end 150 bp (PE150) platform with a minimum coverage of 20 million reads per sample.

#### Data processing/batch correction for bulk RNA seq data

RNA-seq count data were first processed and quality-controlled prior to analysis. One epilepsy-associated line was excluded, and one low-quality sample identified in QC was removed. Genes with zero counts across all samples were filtered out, and data from both experiments were combined. Counts were processed in DESeq2 using its internal median-of-ratios normalization, and a regularized log transformation (rlog) was applied for downstream quality control. Quality assessment included evaluation of library size distributions, sample-to-sample distances, and principal component analysis (PCA). To visualize batch-associated variance, exploratory corrections were performed with both limma’s removeBatchEffect and sva’s ComBat, using experiment date as the batch variable. For all differential expression analyses, batch was modeled as a covariate in the DESeq2 design formula.

#### Transcriptomic age evaluation

RNA-seq count data were preprocessed and normalized as follows. To remove non-expressed genes, only those with at least 10 reads in at least 20% of samples were retained. The filtered data were then subjected to Relative Log Expression (RLE) normalization, log transformation, and YuGene normalization(*103*). Missing expression values for clock genes absent from the dataset were imputed using their respective precomputed average values. The resulting normalized expression profiles were centered to the median profile of DMSO-treated control samples. Transcriptomic age (tAge) for each sample was estimated using composite Elastic Net– based multi-species, multi-tissue transcriptomic clocks of chronological age and expected mortality(*71*). Module-specific transcriptomic clocks of chronological age and expected mortality were applied to the scaled relative expression profiles within the same analytical framework. The resulting tAge estimates from module-specific clocks were standardized following adjustment for patient ID. To compare tAge estimates derived from composite and module-specific clocks between the groups (EL vs non-EL, or DMSO-vs Thapsigargin-treated cells), one-way ANOVA was utilized. The effect of thapsigargin on tAge was evaluated using an ANOVA model with patient ID included as a covariate to account for paired measurements. Resulting p-values were corrected for multiple comparisons using the Benjamini–Hochberg procedure.

#### Gene ontology analysis/heatmap generation

Gene Set Enrichment Analysis was run in preranked mode using the Broad Institute’s GSEA software. Ranked gene lists were generated from DESeq2 differential expression results, ordering genes by the Wald test statistic. Enrichment was assessed against gene sets from the Molecular Signatures Database (MSigDB), with significance determined by permutation testing and false discovery rate (FDR) correction. For heatmap visualization, counts were normalized to counts per million (CPM) and log₂-transformed. Heatmaps were generated from these values for the top genes of interest and curated gene sets.

#### Live cell calcium imaging

Calcium activity was assessed in the control cell lines (4 lines) and in the EL, centenarian cell lines (9 lines) expressing the calcium indicator GCaMP.8s [VB240904-1449tnm]. Each cell line had 4 representative fields of view (FOV) assessed. Time series were captured on the Zeiss LSM 710-Live Duo Confocal at 1 Hz for 180 seconds using a 10X 0.45 aperture objective and incubator attachment (5% CO_2_ and 37 C, PeCon). Each FOV had 15 manually selected regions of interest (ROIs). 15 individual neuron cell body ROIs were selected by hand in each time series and background subtracted based on a measured non-neuronal ROI. The Activity trace for each neuron was calculated as (F-F_o_)/ F_o_ = ΔF/F_o_; the mean fluorescence of the neuron (mean value of the ROI) at each time point, F, normalized by F_o_, the mean of the lowest 1% of measurements for that neuron over the entire trial (*i.e.* the mean of the bottom 1% of measured values). ΔF/F_o_ of measured neurons were plotted on activity heatmaps as in **Figure 3C**. Neuron activation events were detected using a peak prominence of 0.6 in the fluorescence trace (via Peak Prominence peak detection in Matlab)(*104*). A neuron was considered “active” if it had at least one flagged peak during the 180s time series. Width of peak was defined as the time between the onset and end of a calcium activation event, and the interpeak interval was defined as the time between two calcium events in the same neuron. Data was averaged per neuron and findings are summarized in **Figure 3**. Percent active measurements were analyzed using a chi-squared test for independence. WOP and IPI assessments were done using an unpaired two-tailed t-test using Welch’s correction.

#### Comparison of EL and non-EL Thapsigargin response

EL and non-EL neurons were exposed to an acute 500 nM dose of thapsigargin (Fisher Scientific, AC328570050) for 24 hours. Following dosing, RNA was isolated and profiled using bulk RNA sequencing through Novogene Co. as described. Differential expression analysis was performed in DESeq2 using raw counts with a model that included experimental batch, donor group (exceptional longevity versus control), treatment condition (DMSO versus thapsigargin), and an interaction term between donor group and treatment. This interaction term specifically tested whether the transcriptional response to thapsigargin differed between EL and control neurons. DESeq2’s standard workflow was used, including internal normalization for library size, estimation of gene-wise dispersion parameters, and fitting of negative binomial models. Results were extracted for the interaction term, and significance was determined using the Benjamini-Hochberg false discovery rate (FDR).

#### Composite analysis/comparison of EL resiliency signature to LOAD dataset

The EL-stress resiliency signature was derived from the interaction-term differential expression analysis in iPSC-derived neurons, selecting genes with Benjamini-Hochberg adjusted FDR < 0.05. Direction was assigned based on the interaction log₂ fold change: genes with positive interaction log₂ fold change values (greater change in EL neurons relative to controls) were grouped as “upregulated,” while genes with negative values (smaller change in EL neurons relative to controls) were grouped as “downregulated.” A publicly available late-onset Alzheimer’s disease (LOAD) bulk expression dataset (GEO: GSE44772; Zhang et al., Cell 2013), including 129 LOAD patients and 101 controls across dorsolateral prefrontal cortex (DLPFC), visual cortex, and cerebellum (690 samples total), was preprocessed prior to signature projection. Processing steps included retaining probes with mapped gene symbols, collapsing duplicate symbols by selecting the probe with the highest variance, updating feature identifiers to gene symbols, and removing features with missing expression values. Gene set variation analysis (GSVA; R package GSVA) was applied to compute sample-level enrichment scores for the upregulated and downregulated signature sets using a Gaussian kernel. A composite resilience score was then defined as the difference between the upregulated and downregulated GSVA scores (up - down), providing a single quantitative measure of signature activity per sample. Statistical modeling of the composite score was performed using linear mixed-effects models fit with the lme4 package, with estimated marginal means (EMMs) and contrasts obtained using the emmeans package. Fixed effects included LOAD status (LOAD vs control), brain region, and gender, with a random intercept for patient to account for multiple regions per donor. An additive model was compared to one including a LOAD by region interaction using a likelihood-ratio test fit by maximum likelihood (ML), and final estimates were obtained under restricted maximum likelihood (REML). Region-specific EMMs and LOAD-Control differences were derived from the interaction model, while the overall LOAD-Control difference was reported from the additive model to avoid averaging across the interaction.

#### Composite analysis and comparison of EL resiliency signature to TCGA cancer datasets

The EL-stress resiliency signature (upregulated and downregulated gene sets from the iPSC interaction-term analysis) was projected onto TCGA RNA-seq primary tumor datasets obtained via the TCGAbiolinks R package. Within each cancer type, raw counts were processed using DESeq2 median-of-ratios normalization and variance-stabilizing transformation (vst). Gene set variation analysis (GSVA; R package GSVA) was applied to compute sample-level enrichment scores for the upregulated and downregulated signature sets using a Gaussian kernel. A composite resilience score was then defined as the difference between the upregulated and downregulated GSVA scores (up - down). For head and neck squamous cell carcinoma (HNSC), HPV-positive cases were excluded for the survival analyses. Overall survival associations were evaluated using Cox proportional hazards models (survival package) fit separately within each cancer type, including the composite GSVA score (z-scored within cancer) and chronological age as covariates; hazard ratios, 95% confidence intervals, and p-values were extracted. To assess tumor-normal differences, paired tumor and adjacent normal samples (where available) were processed jointly per cancer type by vst-normalizing the combined count matrix in DESeq2 as above, computing GSVA scores as above, and testing paired differences in composite score GSVA with Wilcoxon signed-rank tests. Finally, global Kolmogorov-Smirnov tests were applied across cancers to evaluate whether the distribution of nominal p-values from survival and tumor-normal comparisons was enriched near zero relative to the uniform distribution.

Application of transcriptional clocks: Alex T

#### Mitochondrial characterization using MitoTracker and Seahorse assays

To assess mitochondrial dynamics of produced neurons from EL and non-EL groups, a combination of MitoTracker dyes were employed. MitoTracker Green FM (ThermoFisher, M7514) was used to assess total mitochondrial load and MitoTracker Red CMXRos (ThermoFisher, M7512) was used to assess polarized mitochondria. Cells were stained using 200 nM MitoGreen and MitoRed for 20 minutes at 37C. Following staining, cells were washed and profiled via flow cytometry or imaged using a confocal microscope. Given reported evidence that mitochondrial changes are rapid and short lived following thapsigargin exposure in a biphasic pattern(*76*), cells were exposed to either 500 nM or 1 µM thapsigargin for 5 min before profiling via flow cytometry to capture potential mitochondrial changes (**Supplemental Figure 4**).

A Mito Stress Assay was conducted using a Seahorse XFe96 Instrument. Following optimization it was determined that drug concentrations were optimal at the following concentrations: Oligomycin – 2.5 µM, FCCP – 1.5 µM, Rotenone - 1 µM, Antimycin A – 1 µM. Hoechst staining was conducted to allow for cell number normalization using a plate reader and comparisons were performed using Agilent Seahorse Analytics software.

## Supplemental Figure Legends

**Fig. S1.**
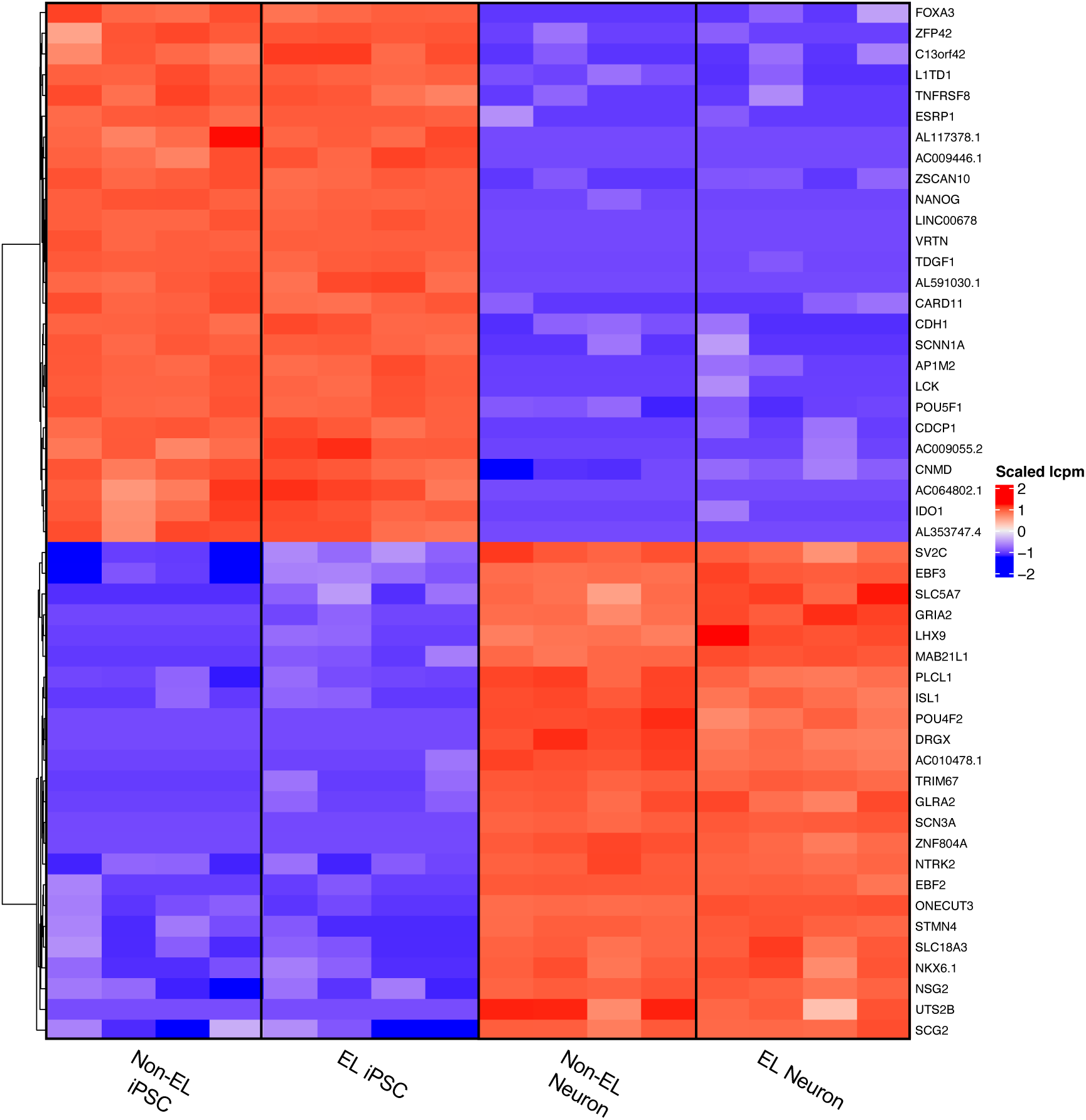
Efficient patterning of forebrain cortical neurons from EL and non-EL iPSCs. A) Heatmap displaying top 50 differentially expressed genes (DEGs, LogFC FDR<0.05) between EL and non-EL (BU) iPSCs and corresponding iPSC-derived neurons (Red = upregulated, Blue = downregulated).

**Fig. S2.**
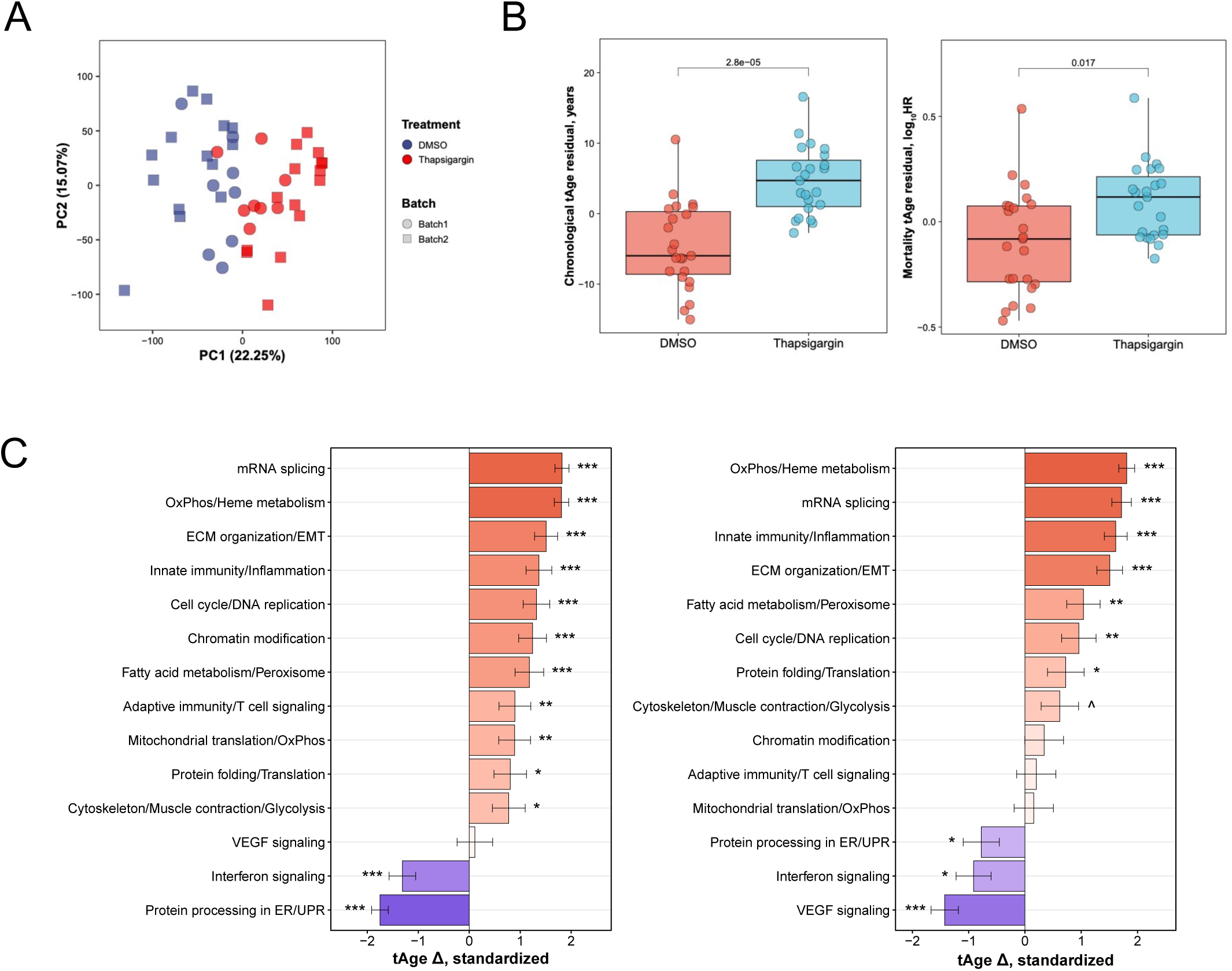
Transcriptomic clock age estimates reveal age increases and stress response pathway enrichment following thapsigargin treatment. A) PCA analysis of transcriptional response between DMSO vehicle (blue) and Thapsigargin treated EL and Non-EL neurons from batches 1 (circle) and 2 (square). B) Multispecies conventional Chronological tAge (left) and mortality tAge (right) predictions of EL and Non-EL neurons treated with DMSO or Thapsigargin (paired statistical test). C) Pathway enrichment following module-specific Chronological tAge (left) and Mortality tAge (right) predictions of EL and Non-EL neurons treated with Thapsigargin compared to DMSO treatment (*=p<0.05, **=p<0.01, ***=p<0.001).

**Fig. S3.**
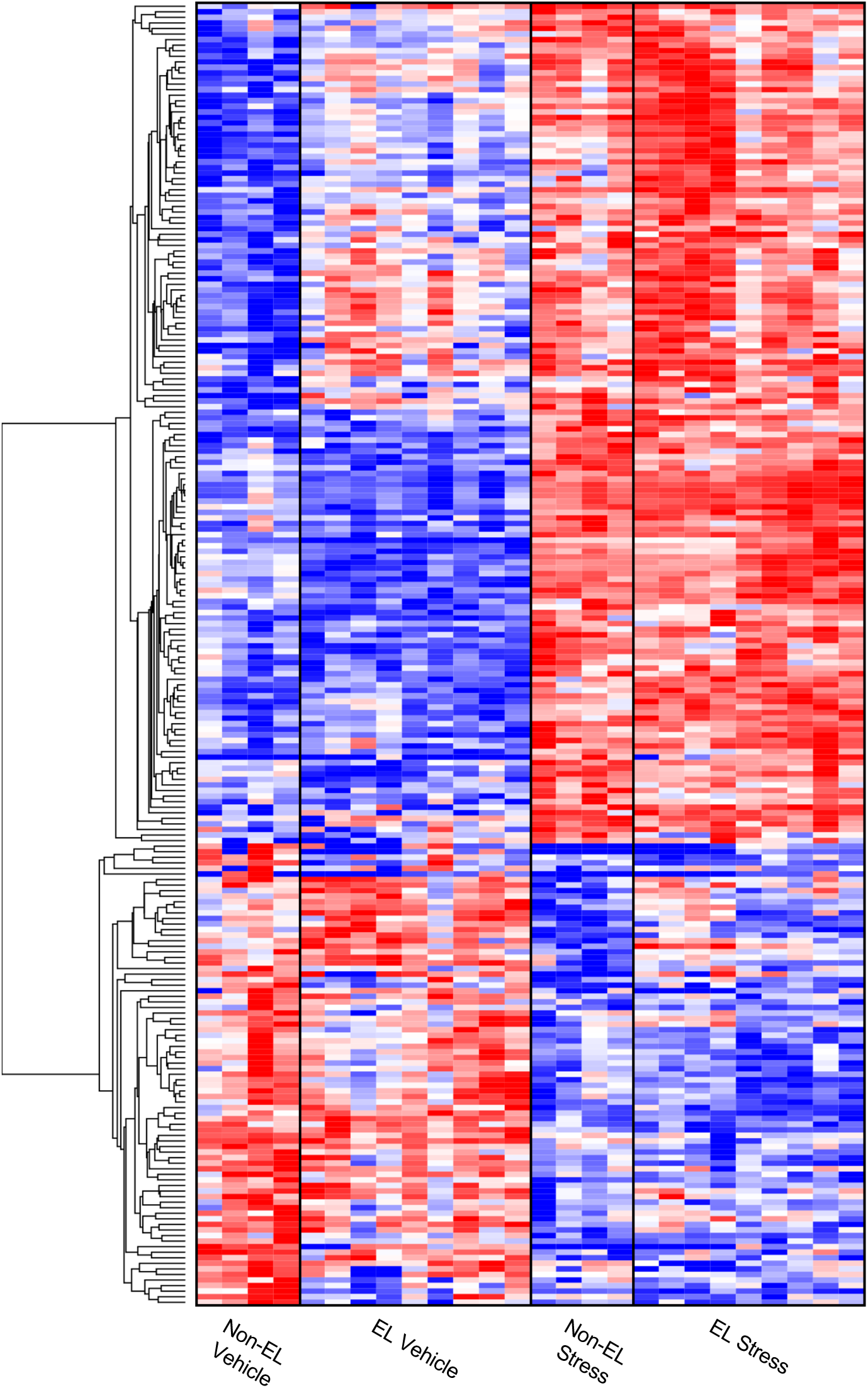
Thapsigargin exposure yields a global molecular response signature in iPSC-derived neurons. A) Heatmap displaying top 200 differentially expressed genes (DEGs) of EL and non-EL neurons (grouped) following global ER stress (Red = upregulated, Blue = downregulated).

**Fig. S4.**
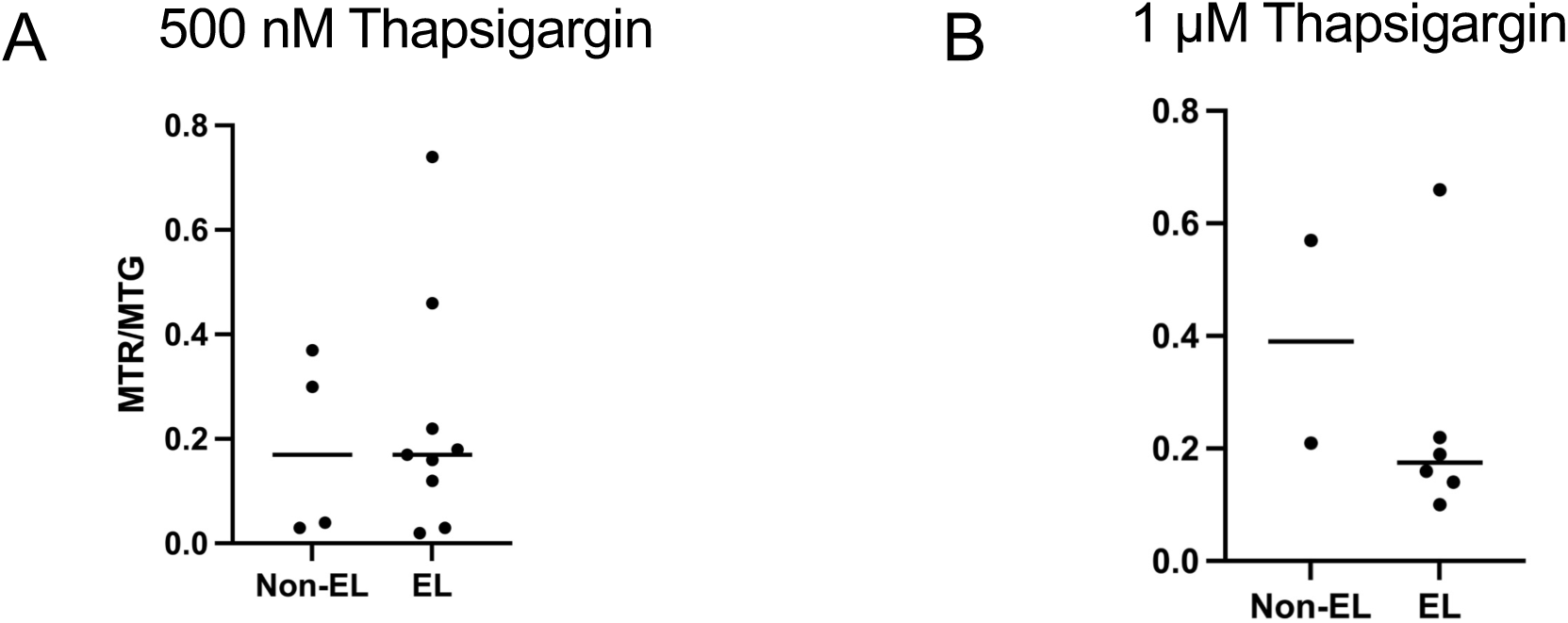
EL neuron mitochondrial dynamics is independent of ER stress response. Quantification of MTR/MTG ratio of 4 non-EL and 9 EL cell lines profiled in flow cytometry MitoTracker experiments following A) 500 nM Thapsigargin exposure or B) 1 µM Thapsigargin exposure for 5 min.

**Movie S1. Representative live calcium imaging of iPSC-derived neurons.**

## References and Notes

1. United Nations, World Population Prospects 2024: Summary of Results (2024).

2. A. Garmany, S. Yamada, A. Terzic, Longevity leap: mind the healthspan gap. npj Regen Med 6, 1–7 (2021).

3. J. Guo, X. Huang, L. Dou, M. Yan, T. Shen, W. Tang, J. Li, Aging and aging-related diseases: from molecular mechanisms to interventions and treatments. Sig Transduct Target Ther 7, 1–40 (2022).

4. A. J. Scott, M. Ellison, D. A. Sinclair, The economic value of targeting aging. Nature Aging 1, 616–623 (2021).

5. C. López-Otín, M. A. Blasco, L. Partridge, M. Serrano, G. Kroemer, The Hallmarks of Aging. Cell 153, 1194–1217 (2013).

6. M.-L. Díaz-Hung, C. Hetz, Proteostasis and resilience: on the interphase between individual’s and intracellular stress. Trends in Endocrinology & Metabolism 33, 305– 317 (2022).

7. G. Hannum, J. Guinney, L. Zhao, L. Zhang, G. Hughes, S. Sadda, B. Klotzle, M. Bibikova, J.-B. Fan, Y. Gao, R. Deconde, M. Chen, I. Rajapakse, S. Friend, T. Ideker, K. Zhang, Genome-wide Methylation Profiles Reveal Quantitative Views of Human Aging Rates. Molecular Cell 49, 359–367 (2013).

8. J.-H. Yang, M. Hayano, P. T. Griffin, J. A. Amorim, M. S. Bonkowski, J. K. Apostolides, E. L. Salfati, M. Blanchette, E. M. Munding, M. Bhakta, Y. C. Chew, W. Guo, X. Yang, S. Maybury-Lewis, X. Tian, J. M. Ross, G. Coppotelli, M. V. Meer, R. Rogers-Hammond, D. L. Vera, Y. R. Lu, J. W. Pippin, M. L. Creswell, Z. Dou, C. Xu, S. J. Mitchell, A. Das, B. L. O’Connell, S. Thakur, A. E. Kane, Q. Su, Y. Mohri, E. K. Nishimura, L. Schaevitz, N. Garg, A.-M. Balta, M. A. Rego, M. Gregory-Ksander, T. C. Jakobs, L. Zhong, H. Wakimoto, J. E. Andari, D. Grimm, R. Mostoslavsky, A. J. Wagers, K. Tsubota, S. J. Bonasera, C. M. Palmeira, J. G. Seidman, C. E. Seidman, N. S. Wolf, J. A. Kreiling, J. M. Sedivy, G. F. Murphy, R. E. Green, B. A. Garcia, S. L. Berger, P. Oberdoerffer, S. J. Shankland, V. N. Gladyshev, B. R. Ksander, A. R. Pfenning, L. A. Rajman, D. A. Sinclair, Loss of epigenetic information as a cause of mammalian aging. Cell 186, 305–326.e27 (2023).

9. L. Hayflick, Entropy Explains Aging, Genetic Determinism Explains Longevity, and Undefined Terminology Explains Misunderstanding Both. PLOS Genetics 3, e220 (2007).

10. K. Mozhui, A. T. Lu, C. Z. Li, A. Haghani, J. V. Sandoval-Sierra, Y. Wu, R. W. Williams, S. Horvath, Genetic Loci and Metabolic States Associated With Murine Epigenetic Aging. bioRxiv, 2021.06.23.449634 (2022).

11. E. Dent, F. C. Martin, H. Bergman, J. Woo, R. Romero-Ortuno, J. D. Walston, Management of frailty: opportunities, challenges, and future directions. The Lancet 394, 1376–1386 (2019).

12. R. S. Vasan, Biomarkers of Cardiovascular Disease. Circulation 113, 2335–2362 (2006).

13. O. Hansson, Biomarkers for neurodegenerative diseases. Nat Med 27, 954–963 (2021).

14. V. K. Sarhadi, G. Armengol, Molecular Biomarkers in Cancer. Biomolecules 12, 1021 (2022).

15. S. J. Olshansky, From Lifespan to Healthspan. JAMA 320, 1323–1324 (2018).

16. S. L. Andersen, Centenarians as Models of Resistance and Resilience to Alzheimer’s Disease and Related Dementias. Adv Geriatr Med Res 2, e200018 (2020).

17. M. A. Islam, U. Sehar, O. F. Sultana, U. Mukherjee, M. Brownell, S. Kshirsagar, P. H. Reddy, SuperAgers and centenarians, dynamics of healthy ageing with cognitive resilience. Mechanisms of Ageing and Development 219, 111936 (2024).

18. R. Hitt, Y. Young-Xu, M. Silver, T. Perls, Centenarians: the older you get, the healthier you have been. The Lancet 354, 652 (1999).

19. J. Evert, E. Lawler, H. Bogan, T. Perls, Morbidity Profiles of Centenarians: Survivors, Delayers, and Escapers. The journals of gerontology. Series A, Biological sciences and medical sciences 58, 232–M237 (2003).

20. D. F. Terry, M. A. Wilcox, M. A. McCormick, T. T. Perls, Cardiovascular Disease Delay in Centenarian Offspring. The journals of gerontology. Series A, Biological sciences and medical sciences 59, 385–M389 (2004).

21. J. F. Fries, Aging, Natural Death, and the Compression of Morbidity. The New England Journal of Medicine 303, 130–135 (1980).

22. T. Perls, Successful aging and its subtypes in centenarians: The Chinese experience. Journal of the American Geriatrics Society n/a.

23. S. L. Andersen, P. Sebastiani, D. A. Dworkis, L. Feldman, T. T. Perls, Health span approximates life span among many supercentenarians: compression of morbidity at the approximate limit of life span. J Gerontol A Biol Sci Med Sci 67, 395–405 (2012).

24. E. A. Schoenhofen, D. F. Wyszynski, S. Andersen, J. Pennington, R. Young, D. F. Terry, T. T. Perls, Characteristics of 32 Supercentenarians. Journal of the American Geriatrics Society 54, 1237–1240 (2006).

25. S. Horvath, C. Pirazzini, M. G. Bacalini, D. Gentilini, A. M. D. Blasio, M. Delledonne, D. Mari, B. Arosio, D. Monti, G. Passarino, F. D. Rango, P. D’Aquila, C. Giuliani, E. Marasco, S. Collino, P. Descombes, P. Garagnani, C. Franceschi, Decreased epigenetic age of PBMCs from Italian semi-supercentenarians and their offspring. Aging 7 (2015).

26. P. Sebastiani, A. Federico, M. Morris, A. Gurinovich, T. Tanaka, K. B. Chandler, S. L. Andersen, G. Denis, C. E. Costello, L. Ferrucci, L. Jennings, D. J. Glass, S. Monti, T. T. Perls, Protein signatures of centenarians and their offspring suggest centenarians age slower than other humans. Aging Cell 20, e13290 (2021).

27. T. T. Karagiannis, T. W. Dowrey, C. Villacorta-Martin, M. Montano, E. Reed, A. C. Belkina, S. L. Andersen, T. T. Perls, S. Monti, G. J. Murphy, P. Sebastiani, Multi-modal profiling of peripheral blood cells across the human lifespan reveals distinct immune cell signatures of aging and longevity. EBioMedicine 90, 104514 (2023).

28. K. Hashimoto, T. Kouno, T. Ikawa, N. Hayatsu, Y. Miyajima, H. Yabukami, T. Terooatea, T. Sasaki, T. Suzuki, M. Valentine, G. Pascarella, Y. Okazaki, H. Suzuki, J. W. Shin, A. Minoda, I. Taniuchi, H. Okano, Y. Arai, N. Hirose, P. Carninci, Single-cell transcriptomics reveals expansion of cytotoxic CD4 T cells in supercentenarians. PNAS Plus 116, 24242–24251 (2019).

29. S. Park, A. Gianotti-Sommer, F. J. Molina-Estevez, K. Vanuytsel, N. Skvir, A. Leung, S. S. Rozelle, E. M. Shaikho, I. Weir, Z. Jiang, H.-Y. Luo, D. H. K. Chui, M. S. Figueiredo, A. Alsultan, A. Al-Ali, P. Sebastiani, M. H. Steinberg, G. Mostoslavsky, G. J. Murphy, A Comprehensive, Ethnically Diverse Library of Sickle Cell Disease-Specific Induced Pluripotent Stem Cells. Stem Cell Reports 8, 1076– 1085 (2017).

30. H. Okano, S. Morimoto, iPSC-based disease modeling and drug discovery in cardinal neurodegenerative disorders. Cell Stem Cell 29, 189–208 (2022).

31. E. R. Adams, V. G. Nolan, S. L. Andersen, T. T. Perls, D. F. Terry, Centenarian offspring: start healthier and stay healthier. J Am Geriatr Soc 56, 2089–2092 (2008).

32. T. T. Perls, E. Bubrick, C. G. Wager, J. Vijg, L. Kruglyak, Siblings of centenarians live longer. The Lancet 351, 1560 (1998).

33. T. T. Perls, J. Wilmoth, R. Levenson, M. Drinkwater, M. Cohen, H. Bogan, E. Joyce, S. Brewster, L. Kunkel, A. Puca, Life-long sustained mortality advantage of siblings of centenarians. Proceedings of the National Academy of Sciences 99, 8442–8447 (2002).

34. T. Perls, D. F. Terry, M. Silver, M. Shea, J. Bowen, E. Joyce, S. B. Ridge, R. Fretts, M. Daly, S. Brewster, A. Puca, L. Kunkel, The Molecular Genetics of Aging. Results and Problems in Cell Differentiation 29, 1–20 (2000).

35. S. Milman, N. Barzilai, K. A. Wilson, O. Van der Willik, S. Lederman, T. Perls, T. Gao, A. M. Leahy, P. Jain, A. Montgomery, A. R. Shuldiner, SuperAger Initiative: unlocking the genetic potential of exceptional longevity. Nat Aging, 1–2 (2023).

36. P. Sebastiani, T. T. Perls, The Genetics of Extreme Longevity: Lessons from the New England Centenarian Study. Frontiers in Genetics 3, 277 (2012).

37. P. Sebastiani, N. Solovieff, A. T. DeWan, K. M. Walsh, A. Puca, S. W. Hartley, E. Melista, S. Andersen, D. A. Dworkis, J. B. Wilk, R. H. Myers, M. H. Steinberg, M. Montano, C. T. Baldwin, J. Hoh, T. T. Perls, Genetic Signatures of Exceptional Longevity in Humans. PLoS ONE 7, e29848 (2012).

38. P. Sebastiani, A. Gurinovich, M. Nygaard, T. Sasaki, B. Sweigart, H. Bae, S. L. Andersen, F. Villa, G. Atzmon, K. Christensen, Y. Arai, N. Barzilai, A. Puca, L. Christiansen, N. Hirose, T. T. Perls, APOE Alleles and Extreme Human Longevity. J Gerontol A Biol Sci Med Sci 74, 44–51 (2019).

39. P. Sebastiani, A. Gurinovich, H. Bae, S. Andersen, A. Malovini, G. Atzmon, F. Villa, A. T. Kraja, D. Ben-Avraham, N. Barzilai, A. Puca, T. T. Perls, Four Genome-Wide Association Studies Identify New Extreme Longevity Variants. J Gerontol A Biol Sci Med Sci 72, 1453–1464 (2017).

40. F. Soldner, D. Hockemeyer, C. Beard, Q. Gao, G. W. Bell, E. G. Cook, G. Hargus, A. Blak, O. Cooper, M. Mitalipova, O. Isacson, R. Jaenisch, Parkinson’s Disease Patient-Derived Induced Pluripotent Stem Cells Free of Viral Reprogramming Factors. Cell 136, 964–977 (2009).

41. G. Cenini, M. Hebisch, V. Iefremova, L. J. Flitsch, Y. Breitkreuz, R. E. Tanzi, D. Y. Kim, M. Peitz, O. Brüstle, Dissecting Alzheimer’s disease pathogenesis in human 2D and 3D models. Molecular and Cellular Neuroscience 110, 103568 (2020).

42. M. Jorfi, C. D’Avanzo, D. Y. Kim, D. Irimia, Three-Dimensional Models of the Human Brain Development and Diseases. Advanced Healthcare Materials 7, 1700723 (2018).

43. M. Jorfi, C. D’Avanzo, R. E. Tanzi, D. Y. Kim, D. Irimia, Human Neurospheroid Arrays for In Vitro Studies of Alzheimer’s Disease. Scientific Reports 8, 2450 (2018).

44. J. Park, I. Wetzel, I. Marriott, D. Dréau, C. D’Avanzo, D. Y. Kim, R. E. Tanzi, H. Cho, A 3D human triculture system modeling neurodegeneration and neuroinflammation in Alzheimer’s disease. Nat Neurosci 21, 941–951 (2018).

45. L. Quinti, J. Park, E. Brand, R. E. Tanzi, D. Y. Kim, Neurotherapeutics in the Era of Translational Medicine. 311–331 (2021).

46. R. M. Giadone, D. C. Liberti, T. M. Matte, J. D. Rosarda, C. Torres-Arancivia, S. Ghosh, J. K. Diedrich, S. Pankow, N. Skvir, J. C. Jean, J. R. Yates, A. A. Wilson, L. H. Connors, D. N. Kotton, R. L. Wiseman, G. J. Murphy, Expression of Amyloidogenic Transthyretin Drives Hepatic Proteostasis Remodeling in an Induced Pluripotent Stem Cell Model of Systemic Amyloid Disease. Stem Cell Reports 15, 515–528 (2020).

47. S. Ghosh, C. Villacorta-Martin, J. Lindstrom-Vautrin, D. Kenney, C. S. Golden, C. V. Edwards, V. Sanchorawala, L. H. Connors, R. M. Giadone, G. J. Murphy, Mapping cellular response to destabilized transthyretin reveals cell-and amyloidogenic protein-specific signatures. Amyloid 0, 1–15 (2023).

48. A. Leung, S. K. Nah, W. Reid, A. Ebata, C. M. Koch, S. Monti, J. C. Genereux, R. L. Wiseman, B. Wolozin, L. H. Connors, J. L. Berk, D. C. Seldin, G. Mostoslavsky, D. N. Kotton, G. J. Murphy, Induced Pluripotent Stem Cell Modeling of Multisystemic, Hereditary Transthyretin Amyloidosis. Stem Cell Reports 1, 451–463 (2013).

49. T. W. Dowrey, S. F. Cranston, N. Skvir, Y. Lok, B. Gould, B. Petrowitz, D. Villar, J. Shan, M. James, M. Dodge, A. C. Belkina, R. M. Giadone, S. Milman, P. Sebastiani, T. T. Perls, S. L. Andersen, G. J. Murphy, A longevity-specific bank of induced pluripotent stem cells from centenarians and their offspring. Aging Cell n/a, e14351.

50. F. I. Mahoney, D. W. Barthel, Functional Evaluation: The Barthel Index. Md State Med J 14, 61–65 (1965).

51. A. T. Lu, A. Quach, J. G. Wilson, A. P. Reiner, A. Aviv, K. Raj, L. Hou, A. A. Baccarelli, Y. Li, J. D. Stewart, E. A. Whitsel, T. L. Assimes, L. Ferrucci, S. Horvath, DNA methylation GrimAge strongly predicts lifespan and healthspan. Aging 11, 303–327 (2019).

52. A. T. Higgins-Chen, K. L. Thrush, Y. Wang, C. J. Minteer, P.-L. Kuo, M. Wang, P. Niimi, G. Sturm, J. Lin, A. Z. Moore, S. Bandinelli, C. H. Vinkers, E. Vermetten, B. P. F. Rutten, E. Geuze, C. Okhuijsen-Pfeifer, M. Z. van der Horst, S. Schreiter, S. Gutwinski, J. J. Luykx, M. Picard, L. Ferrucci, E. M. Crimmins, M. P. Boks, S. Hägg, T. T. Hu-Seliger, M. E. Levine, A computational solution for bolstering reliability of epigenetic clocks: implications for clinical trials and longitudinal tracking. Nat Aging 2, 644–661 (2022).

53. M. Fuentealba, L. Rouch, S. Guyonnet, J.-M. Lemaitre, P. de Souto Barreto, B. Vellas, S. Andrieu, D. Furman, A blood-based epigenetic clock for intrinsic capacity predicts mortality and is associated with clinical, immunological and lifestyle factors. Nat Aging 5, 1207–1216 (2025).

54. K. Ying, H. Liu, A. E. Tarkhov, M. C. Sadler, A. T. Lu, M. Moqri, S. Horvath, Z. Kutalik, X. Shen, V. N. Gladyshev, Causality-enriched epigenetic age uncouples damage and adaptation. Nat Aging, 1–16 (2024).

55. S. Horvath, DNA methylation age of human tissues and cell types. Genome biology 14, R115 (2013).

56. M. E. Levine, A. T. Lu, A. Quach, B. H. Chen, T. L. Assimes, S. Bandinelli, L. Hou, A. A. Baccarelli, J. D. Stewart, Y. Li, E. A. Whitsel, J. G. Wilson, A. P. Reiner, A. Aviv, K. Lohman, Y. Liu, L. Ferrucci, S. Horvath, An epigenetic biomarker of aging for lifespan and healthspan. Aging 10, 573–591 (2018).

57. A. Upadhyay, “Neurocalcin delta (NCALD) knockout impairs adult neurogenesis whereas half reduction is a safe therapeutic option for spinal muscular atrophy,” thesis, Universität zu Köln (2019).

58. W. Wang, Z. Zhou, W. Zhao, Y. Huang, R. Tang, K. Ying, Y. Xie, Y. Mao, Molecular cloning, mapping and characterization of the human neurocalcin delta gene (NCALD)1. Biochimica et Biophysica Acta (BBA) - Gene Structure and Expression 1518, 162–167 (2001).

59. M. Riessland, A. Kaczmarek, S. Schneider, K. J. Swoboda, H. Löhr, C. Bradler, V. Grysko, M. Dimitriadi, S. Hosseinibarkooie, L. Torres-Benito, M. Peters, A. Upadhyay, N. Biglari, S. Kröber, I. Hölker, L. Garbes, C. Gilissen, A. Hoischen, G. Nürnberg, P. Nürnberg, M. Walter, F. Rigo, C. F. Bennett, M. J. Kye, A. C. Hart, M. Hammerschmidt, P. Kloppenburg, B. Wirth, Neurocalcin Delta Suppression Protects against Spinal Muscular Atrophy in Humans and across Species by Restoring Impaired Endocytosis. The American Journal of Human Genetics 100, 297–315 (2017).

60. D. Mornet, A. Bonet-Kerrache, Neurocalcin–actin interaction. Biochimica et Biophysica Acta (BBA) - Protein Structure and Molecular Enzymology 1549, 197–203 (2001).

61. R. D. Burgoyne, Neuronal calcium sensor proteins: generating diversity in neuronal Ca2+ signalling. Nat Rev Neurosci 8, 182–193 (2007).

62. S.-Q. Zhang, T. Jiang, M. Li, X. Zhang, Y.-Q. Ren, S.-C. Wei, L.-D. Sun, H. Cheng, Y. Li, X.-Y. Yin, Z.-M. Hu, Z.-Y. Wang, Y. Liu, B.-R. Guo, H.-Y. Tang, X.-F. Tang, Y.-T. Ding, J.-B. Wang, P. Li, B.-Y. Wu, W. Wang, X.-F. Yuan, J.-S. Hou, W.-W. Ha, W.-J. Wang, Y.-J. Zhai, J. Wang, F.-F. Qian, F.-S. Zhou, G. Chen, X.-B. Zuo, X.-D. Zheng, Y.-J. Sheng, J.-P. Gao, B. Liang, P. Li, J. Zhu, F.-L. Xiao, P.-G. Wang, Y. Cui, H. Li, S.-X. Liu, M. Gao, X. Fan, S.-K. Shen, M. Zeng, G.-Q. Sun, Y. Xu, J.-C. Hu, T.-T. He, Y.-R. Li, H.-M. Yang, J. Wang, Z.-Y. Yu, H.-F. Zhang, X. Hu, K. Yang, J. Wang, S.-X. Zhao, Y.-W. Zhou, J.-J. Liu, W.-D. Du, L. Zhang, K. Xia, S. Yang, J. Wang, X.-J. Zhang, Exome sequencing identifies MVK mutations in disseminated superficial actinic porokeratosis. Nat Genet 44, 1156–1160 (2012).

63. D. H. Mauch, K. Nägler, S. Schumacher, C. Göritz, E.-C. Müller, A. Otto, F. W. Pfrieger, CNS Synaptogenesis Promoted by Glia-Derived Cholesterol. Science 294, 1354–1357 (2001).

64. Essential role for cholesterol in synaptic plasticity and neuronal degeneration - Koudinov - 2001 - The FASEB Journal - Wiley Online Library. https://faseb.onlinelibrary.wiley.com/doi/abs/10.1096/fj.00-0815fje.

65. T. C. Harned, R. V. Stan, Z. Cao, R. Chakrabarti, H. N. Higgs, C. C. Y. Chang, T. Y. Chang, Acute ACAT1/SOAT1 Blockade Increases MAM Cholesterol and Strengthens ER-Mitochondria Connectivity. Int J Mol Sci 24, 5525 (2023).

66. M. Fujimoto, T. Hayashi, T.-P. Su, The role of cholesterol in the association of endoplasmic reticulum membranes with mitochondria. Biochem Biophys Res Commun 417, 635–639 (2012).

67. C.-H. Huang, Y.-R. Chu, Y. Ye, X. Chen, Role of HERP and a HERP-related Protein in HRD1-dependent Protein Degradation at the Endoplasmic Reticulum*. Journal of Biological Chemistry 289, 4444–4454 (2014).

68. L. Zhao, C. Rosales, K. Seburn, D. Ron, S. L. Ackerman, Alteration of the unfolded protein response modifies neurodegeneration in a mouse model of Marinesco–Sjögren syndrome. Hum Mol Genet 19, 25–35 (2010).

69. H. Yoshida, T. Matsui, A. Yamamoto, T. Okada, K. Mori, XBP1 mRNA Is Induced by ATF6 and Spliced by IRE1 in Response to ER Stress to Produce a Highly Active Transcription Factor. Cell 107, 881–891 (2001).

70. R. C. Taylor, A. Dillin, XBP-1 is a cell-nonautonomous regulator of stress resistance and longevity. Cell 153, 1435–1447 (2013).

71. A. Tyshkovskiy, D. Kholdina, K. Ying, M. Davitadze, A. Molière, Y. Tongu, T. Kasahara, L. M. Kats, A. Vladimirova, A. Moldakozhayev, H. Liu, B. Zhang, U. Khasanova, M. Moqri, J. M. V. Raamsdonk, D. E. Harrison, R. Strong, T. Abe, S. E. Dmitriev, V. N. Gladyshev, Transcriptomic Hallmarks of Mortality Reveal Universal and Specific Mechanisms of Aging, Chronic Disease, and Rejuvenation. bioRxiv [Preprint] (2024). 10.1101/2024.07.04.601982.

72. C. López-Otín, M. A. Blasco, L. Partridge, M. Serrano, G. Kroemer, Hallmarks of aging: An expanding universe. Cell, doi: 10.1016/j.cell.2022.11.001.

73. Md. T. Islam, Oxidative stress and mitochondrial dysfunction-linked neurodegenerative disorders. Neurological Research 39, 73–82 (2017).

74. M. Y. Vyssokikh, S. Holtze, O. A. Averina, K. G. Lyamzaev, A. A. Panteleeva, M. V. Marey, R. A. Zinovkin, F. F. Severin, M. V. Skulachev, N. Fasel, T. B. Hildebrandt, V. P. Skulachev, Mild depolarization of the inner mitochondrial membrane is a crucial component of an anti-aging program. Proceedings of the National Academy of Sciences 117, 6491–6501 (2020).

75. J. Lytton, M. Westlin, M. R. Hanley, Thapsigargin inhibits the sarcoplasmic or endoplasmic reticulum Ca-ATPase family of calcium pumps. J Biol Chem 266, 17067–17071 (1991).

76. J. R. Hom, J. S. Gewandter, L. Michael, S.-S. Sheu, Y. Yoon, Thapsigargin induces biphasic fragmentation of mitochondria through calcium-mediated mitochondrial fission and apoptosis. Journal of Cellular Physiology 212, 498–508 (2007).

77. T. D. Burton, A. O. Fedele, J. Xie, L. Y. Sandeman, C. G. Proud, The gene for the lysosomal protein LAMP3 is a direct target of the transcription factor ATF4. J Biol Chem 295, 7418–7430 (2020).

78. H. Mujcic, T. Rzymski, K. M. A. Rouschop, M. Koritzinsky, M. Milani, A. L. Harris, B. G. Wouters, Hypoxic activation of the unfolded protein response (UPR) induces expression of the metastasis-associated gene LAMP3. Radiotherapy and Oncology 92, 450–459 (2009).

79. W. Ma, B. Ortiz-Quintero, R. Rangel, M. R. McKeller, S. Herrera-Rodriguez, E. F. Castillo, K. S. Schluns, M. Hall, H. Zhang, W.-K. Suh, H. Okada, T. W. Mak, Y. Zhou, M. R. Blackburn, H. Martinez-Valdez, Coordinate activation of inflammatory gene networks, alveolar destruction and neonatal death in AKNA deficient mice. Cell Res 21, 1564–1577 (2011).

80. B. Zhang, C. Gaiteri, L.-G. Bodea, Z. Wang, J. McElwee, A. A. Podtelezhnikov, C. Zhang, T. Xie, L. Tran, R. Dobrin, E. Fluder, B. Clurman, S. Melquist, M. Narayanan, C. Suver, H. Shah, M. Mahajan, T. Gillis, J. Mysore, M. E. MacDonald, J. R. Lamb, D. A. Bennett, C. Molony, D. J. Stone, V. Gudnason, A. J. Myers, E. E. Schadt, H. Neumann, J. Zhu, V. Emilsson, Integrated Systems Approach Identifies Genetic Nodes and Networks in Late-Onset Alzheimer’s Disease. Cell 153, 707–720 (2013).

81. The Cancer Genome Atlas Pan-Cancer analysis project | Nature Genetics. https://www.nature.com/articles/ng.2764.

82. D. Simkin, E. Kiskinis, Modeling Pediatric Epilepsy Through iPSC-Based Technologies. Epilepsy Curr 18, 240–245 (2018).

83. S. S. Korshunov, V. P. Skulachev, A. A. Starkov, High protonic potential actuates a mechanism of production of reactive oxygen species in mitochondria. FEBS Lett 416, 15–18 (1997).

84. S. Miwa, M. D. Brand, Mitochondrial matrix reactive oxygen species production is very sensitive to mild uncoupling. Biochem Soc Trans 31, 1300–1301 (2003).

85. M. Keane, J. Semeiks, A. E. Webb, Y. I. Li, V. Quesada, T. Craig, L. B. Madsen, S. van Dam, D. Brawand, P. I. Marques, P. Michalak, L. Kang, J. Bhak, H.-S. Yim, N. V. Grishin, N. H. Nielsen, M. P. Heide-Jørgensen, E. M. Oziolor, C. W. Matson, G. M. Church, G. W. Stuart, J. C. Patton, J. C. George, R. Suydam, K. Larsen, C. López-Otín, M. J. O’Connell, J. W. Bickham, B. Thomsen, J. P. de Magalhães, Insights into the evolution of longevity from the bowhead whale genome. Cell Rep 10, 112–122 (2015).

86. R. Buffenstein, Negligible senescence in the longest living rodent, the naked mole-rat: insights from a successfully aging species. J Comp Physiol B 178, 439– 445 (2008).

87. E. B. Kim, X. Fang, A. A. Fushan, Z. Huang, A. V. Lobanov, L. Han, S. M. Marino, X. Sun, A. A. Turanov, P. Yang, S. H. Yim, X. Zhao, M. V. Kasaikina, N. Stoletzki, C. Peng, P. Polak, Z. Xiong, A. Kiezun, Y. Zhu, Y. Chen, G. V. Kryukov, Q. Zhang, L. Peshkin, L. Yang, R. T. Bronson, R. Buffenstein, B. Wang, C. Han, Q. Li, L. Chen, W. Zhao, S. R. Sunyaev, T. J. Park, G. Zhang, J. Wang, V. N. Gladyshev, Genome sequencing reveals insights into physiology and longevity of the naked mole rat. Nature 479, 223–227 (2011).

88. R. Buffenstein, J. U. M. Jarvis, The naked mole rat--a new record for the oldest living rodent. Sci Aging Knowledge Environ 2002, pe7 (2002).

89. X. Tian, J. Azpurua, C. Hine, A. Vaidya, M. Myakishev-Rempel, J. Ablaeva, Z. Mao, E. Nevo, V. Gorbunova, A. Seluanov, High-molecular-mass hyaluronan mediates the cancer resistance of the naked mole rat. Nature 499, 346–349 (2013).

90. J. Azpurua, Z. Ke, I. X. Chen, Q. Zhang, D. N. Ermolenko, Z. D. Zhang, V. Gorbunova, A. Seluanov, Naked mole-rat has increased translational fidelity compared with the mouse, as well as a unique 28S ribosomal RNA cleavage. Proc Natl Acad Sci U S A 110, 17350–17355 (2013).

91. I. Heinze, M. Bens, E. Calzia, S. Holtze, O. Dakhovnik, A. Sahm, J. M. Kirkpatrick, K. Szafranski, N. Romanov, S. N. Sama, K. Holzer, S. Singer, M. Ermolaeva, M. Platzer, T. Hildebrandt, A. Ori, Species comparison of liver proteomes reveals links to naked mole-rat longevity and human aging. BMC Biol 16, 82 (2018).

92. K. A. Rodriguez, P. A. Osmulski, A. Pierce, S. T. Weintraub, M. Gaczynska, R. Buffenstein, A cytosolic protein factor from the naked mole-rat activates proteasomes of other species and protects these from inhibition. Biochim Biophys Acta 1842, 2060–2072 (2014).

93. M. Moqri, C. Herzog, J. R. Poganik, K. Ying, J. N. Justice, D. W. Belsky, A. T. Higgins-Chen, B. H. Chen, A. A. Cohen, G. Fuellen, S. Hägg, R. E. Marioni, M. Widschwendter, K. Fortney, P. O. Fedichev, A. Zhavoronkov, N. Barzilai, J. Lasky-Su, D. P. Kiel, B. K. Kennedy, S. Cummings, P. E. Slagboom, E. Verdin, A. B. Maier, V. Sebastiano, M. P. Snyder, V. N. Gladyshev, S. Horvath, L. Ferrucci, Validation of biomarkers of aging. Nat Med 30, 360–372 (2024).

94. M. Moqri, C. Herzog, J. R. Poganik, J. Justice, D. W. Belsky, A. Higgins-Chen, A. Moskalev, G. Fuellen, A. A. Cohen, I. Bautmans, M. Widschwendter, J. Ding, A. Fleming, J. Mannick, J.-D. J. Han, A. Zhavoronkov, N. Barzilai, M. Kaeberlein, S. Cummings, B. K. Kennedy, L. Ferrucci, S. Horvath, E. Verdin, A. B. Maier, M. P. Snyder, V. Sebastiano, V. N. Gladyshev, Biomarkers of aging for the identification and evaluation of longevity interventions. Cell 186, 3758–3775 (2023).

95. C. M. S. Herzog, L. J. E. Goeminne, J. R. Poganik, N. Barzilai, D. W. Belsky, J. Betts-LaCroix, B. H. Chen, M. Chen, A. A. Cohen, S. R. Cummings, P. O. Fedichev, L. Ferrucci, A. Fleming, K. Fortney, D. Furman, V. Gorbunova, A. Higgins-Chen, L. Hood, S. Horvath, J. N. Justice, D. P. Kiel, G. A. Kuchel, J. Lasky-Su, N. K. LeBrasseur, A. B. Maier, B. Schilling, V. Sebastiano, P. E. Slagboom, M. P. Snyder, E. Verdin, M. Widschwendter, A. Zhavoronkov, M. Moqri, V. N. Gladyshev, Challenges and recommendations for the translation of biomarkers of aging. Nat Aging 4, 1372–1383 (2024).

96. R. C. M. Hennekam, Pathophysiology of premature aging characteristics in Mendelian progeroid disorders. European Journal of Medical Genetics 63, 104028 (2020).

97. S. Horvath, J. Oshima, G. M. Martin, A. T. Lu, A. Quach, H. Cohen, S. Felton, M. Matsuyama, D. Lowe, S. Kabacik, J. G. Wilson, A. P. Reiner, A. Maierhofer, J. Flunkert, A. Aviv, L. Hou, A. A. Baccarelli, Y. Li, J. D. Stewart, E. A. Whitsel, L. Ferrucci, S. Matsuyama, K. Raj, Epigenetic clock for skin and blood cells applied to Hutchinson Gilford Progeria Syndrome and ex vivo studies. Aging (Albany NY) 10, 1758–1775 (2018).

98. G. Monnerat, T. H. Kasai-Brunswick, K. D. Asensi, D. Silva Dos Santos, R. A. Q. Barbosa, F. Cristina Paccola Mesquita, J. P. Calvancanti Albuquerque, P. F. Raphaela, C. Wendt, K. Miranda, G. B. Domont, F. C. S. Nogueira, A. Bastos Carvalho, A. C. Campos de Carvalho, Modelling premature cardiac aging with induced pluripotent stem cells from a hutchinson-gilford Progeria Syndrome patient. Front Physiol 13, 1007418 (2022).

99. N. Daily, J. Elson, T. Wakatsuki, Aging Model for Analyzing Drug-Induced Proarrhythmia Risks Using Cardiomyocytes Differentiated from Progeria-Patient-Derived Induced Pluripotent Stem Cells. Int J Mol Sci 24, 11959 (2023).

100. L. Chen, B. Ge, F. P. Casale, L. Vasquez, T. Kwan, D. Garrido-Martín, S. Watt, Y. Yan, K. Kundu, S. Ecker, A. Datta, D. Richardson, F. Burden, D. Mead, A. L. Mann, J. M. Fernandez, S. Rowlston, S. P. Wilder, S. Farrow, X. Shao, J. J. Lambourne, A. Redensek, C. A. Albers, V. Amstislavskiy, S. Ashford, K. Berentsen, L. Bomba, G. Bourque, D. Bujold, S. Busche, M. Caron, S.-H. Chen, W. Cheung, O. Delaneau, E. T. Dermitzakis, H. Elding, I. Colgiu, F. O. Bagger, P. Flicek, E. Habibi, V. Iotchkova, E. Janssen-Megens, B. Kim, H. Lehrach, E. Lowy, A. Mandoli, F. Matarese, M. T. Maurano, J. A. Morris, V. Pancaldi, F. Pourfarzad, K. Rehnstrom, A. Rendon, T. Risch, N. Sharifi, M.-M. Simon, M. Sultan, A. Valencia, K. Walter, S.-Y. Wang, M. Frontini, S. E. Antonarakis, L. Clarke, M.-L. Yaspo, S. Beck, R. Guigo, D. Rico, J. H. A. Martens, W. H. Ouwehand, T. W. Kuijpers, D. S. Paul, H. G. Stunnenberg, O. Stegle, K. Downes, T. Pastinen, N. Soranzo, Genetic Drivers of Epigenetic and Transcriptional Variation in Human Immune Cells. Cell 167, 1398–1414.e24 (2016).

101. M. J. Aryee, A. E. Jaffe, H. Corrada-Bravo, C. Ladd-Acosta, A. P. Feinberg, K. D. Hansen, R. A. Irizarry, Minfi: a flexible and comprehensive Bioconductor package for the analysis of Infinium DNA methylation microarrays. Bioinformatics 30, 1363–1369 (2014).

102. W. Zhou, T. J. Triche, P. W. Laird, H. Shen, SeSAMe: reducing artifactual detection of DNA methylation by Infinium BeadChips in genomic deletions. Nucleic Acids Research 46, gky691-(2018).

103. K.-A. Lê Cao, F. Rohart, L. McHugh, O. Korn, C. A. Wells, YuGene: a simple approach to scale gene expression data derived from different platforms for integrated analyses. Genomics 103, 239–251 (2014).

104. Y. Lee, M. T. Kim, G. Rhodes, K. Sack, S. J. Son, C. B. Rich, V. B. Kolachalama, C. V. Gabel, V. Trinkaus-Randall, Sustained Ca2+ mobilizations: A quantitative approach to predict their importance in cell-cell communication and wound healing. PLOS ONE 14, e0213422 (2019).

